# Dethroning the Fano Factor: a flexible, model-based approach to partitioning neural variability

**DOI:** 10.1101/165670

**Authors:** Adam S. Charles, Mijung Park, J. Patrick Weller, Gregory D. Horwitz, Jonathan W. Pillow

## Abstract

Neurons in many brain areas exhibit high trial-to-trial variability, with spike counts that are over-dispersed relative to a Poisson distribution. Recent work (Goris et al., 2014) has proposed to explain this variability in terms of a multiplicative interaction between a stochastic gain variable and a stimulus-dependent Poisson firing rate, which produces quadratic relationships between spike count mean and variance. Here we examine this quadratic assumption and propose a more flexible family of models that can account for a more diverse set of mean-variance relationships. Our model contains additive Gaussian noise that is transformed nonlinearly to produce a Poisson spike rate. Different choices of the nonlinear function can give rise to qualitatively different mean-variance relationships, ranging from sub-linear to linear to multiplicative. Intriguingly, a rectified squaring nonlinearity produces a linear mean-variance function, corresponding to responses with constant Fano factor. We describe a computationally efficient method for fitting this model to data, and demonstrate that a majority of neurons in a V1 population are better described by a model with non-quadratic relationship between mean and variance. Lastly, we develop an application to Bayesian adaptive stimulus selection in closed-loop neurophysiology experiments, which shows that accounting for overdispersion can lead to dramatic improvements in adaptive tuning curve estimation.

## 1 Introduction

Quantifying neural variability is crucial for understanding how neurons process and transmit information. This has motivated a large body of work that seeks to characterize the signal and noise governing neural responses to sensory stimuli. A simple but popular approach to this problem models neural spike counts using the Poisson distribution. Under this model, spike count mean and variance are equal. Deviations from this relationship are often characterized by the Fano factor, defined as the ratio of the spike-count variance to the mean (Geisler & Albrecht, 1997; Eden & Kramer, 2010; Shadlen & Newsome, 1998). The Fano factor for a Poisson neuron is therefore equal to one. A Fano factor less than one indicates sub-Poisson variability, a condition referred to as “under-dispersion”; a Fano factor greater than one, on the other hand, indicates greater-than-Poisson variability, commonly known as “over-dispersion”. A substantial literature has shown that Fano factors in a variety of different brain areas differ substantially from 1 (Shadlen & Newsome, 1998; Gur et al., 1997; Barberini et al., 2001; Baddeley et al., 1997; Gershon et al., 1998). Recent work has shown that, not only do neural responses not obey a Poisson distribution, the overall mean-variance relationship is often not well characterized by a line, meaning the data are not consistent with a single Fano factor (Pillow & Scott, 2012b; Gao et al., 2015; Stevenson, 2016; Goris et al., 2014; Wiener & Richmond, 2003; Moshitch & Nelken, 2014). Rather, the spike count variance changes nonlinearly with mean.

Here we develop a flexible model for over-dispersed spike count data that captures a variety of different nonlinear mean-variance relationships. Our approach extends recent work from Goris et al. (2014), which described over-dispersed spike responses in the early visual pathway using a Poisson model with multiplicative noise affecting the trial-to-trial spike rate. This produces a quadratic relationship between mean and variance, so Fano factor increases monotonically with mean spike count.

By contrast, our model seeks to describe a more diverse range of mean-variance relationships while maintaining tractability for simulation and fitting. Our model, which we refer to as the *flexible over-dispersion model*, consists of a stimulus-dependent term, additive Gaussian noise, a point nonlinearity, and conditionally Poisson spiking (see Fig. 1). This framework includes the multiplicative Goris model as a special case when the nonlinearity is exponential, but includes the flexibility to exhibit other behaviors as well: for example, Fano factors that decrease with increasing spike rate, which arises for a rectified-linear nonlinearity, and constant slope linear relationships, corresponding to a constant Fano factor, which arises for a rectified squaring nonlinearity. We show that the model can be tractably fit to data using the Laplace approximation to compute likelihoods, and requires fewer parameters than other models for non-Poisson spike count data such as the modulated binomial or generalized count models (Gao et al., 2015; Stevenson, 2016). We apply our model to the V1 dataset presented in Goris et al. (2014) and show that a rectified power-law nonlinearity provides a better description of most individual neurons than a purely multiplicative model.

**Figure 1:**
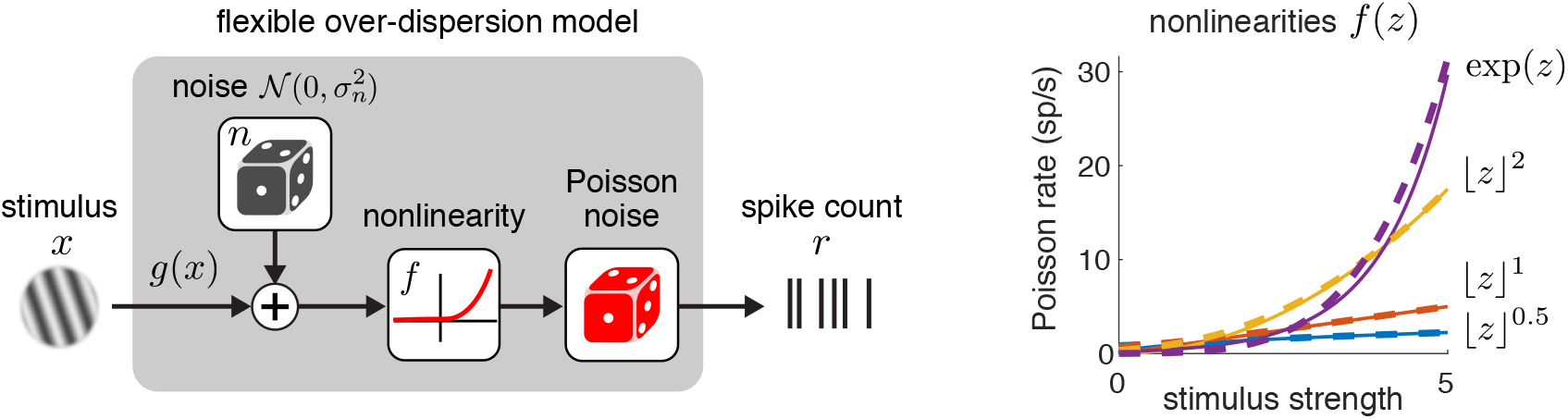
Illustration of the flexible over-dispersion model. **Left**: Model diagram. A stimulus dependent term *g*(**x**) plus a latent zero-mean Gaussian random variable with variance *

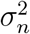

* is transformed by a rectifying nonlinearity *f* (·), whose output drives Poisson spiking. **Right:** The nonlinearity controls the relationship between the stimulus strength and firing rate, and allows for different forms of over-dispersion. Example nonlinearities include hard-rectification raised to a power and the exponential function, which can all be approximated with soft-rectification function raised to a power (eq. 13). Soft-rectification versions are shown as dashed lines, with the power *p* = 3 used to approximate the exponential nonlinearity.

Lastly, we develop an application to adaptive stimulus selection in closed-loop experiments, also known as “active learning”, for characterizing multi-dimensional tuning curves. These methods seek to minimize the amount of time required to estimate tuning curves by selecting stimuli that are as useful or informative as possible about the tuning curve. We use the flexible over-dispersion model in place of the standard Poisson model and demonstrate a marked improvement for tuning curve estimation, both for simulated experiments, as well as for color-tuning maps in awake fixating monkeys.

## 2 Background: Fano factor and the modulated Poisson model

A popular model of neural responses uses the Poisson distribution to describe stimulus-evoked spike counts (Brillinger, 1988; Chichilnisky, 2001; Simoncelli et al., 2004; Paninski, 2004). The basic model specifies a stimulus-dependent spike rate *λ*(**x**) that drives Poisson firing. If we express *λ*(**x**) in units of spikes/bin, the probability of observing *r* spikes in a bin is given by:

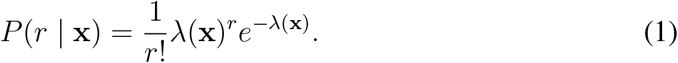

This model makes the strong assumption that the mean and variance are equal: var[*r*] = 𝔼[*r*] = *λ*(**x**), for any stimulus **x**. A sizeable literature has shown assumption is often inaccurate, as spike counts in many brain areas exhibit over-dispersion relative to the Poisson distribution, meaning that variance exceeds the mean (Shadlen & Newsome, 1998; Gur et al., 1997; Barberini et al., 2001; Pillow & Scott, 2012b; Gao et al., 2015; Goris et al., 2014; Stevenson, 2016; Baddeley et al., 1997; Gershon et al., 1998; Tolhurst et al., 1983; Buracas et al., 1998; Carandini, 2004).

A common approach to the phenomenon of over-dispersion is to regard the Fano factor, *F* = var[*r*]*/*𝔼[*r*], as a constant that characterizes the generic degree of overdispersion in neural firing (Geisler & Albrecht, 1997; Shadlen & Newsome, 1998). However, this description is adequate only if spike count variance scales linearly with the mean. Recent work has shown that spike responses in four different early visual areas exhibit variance that grows super-linearly with the mean, meaning that overdispersion (and the Fano factor) increase with the firing rate, inconsistent with a single Fano factor (Goris et al., 2014). Moreover, the Fano factor falls short of providing a complete description of neural spike count statistics, as it does not specify a full probability distribution over spike counts. Such a description is necessary for quantifying information in neural population codes. In light of these shortcomings, we feel that the Fano factor should be set aside as the default statistic for characterizing neural overdispersion; rather, researchers should consider the full curve describing how variance changes as a function of mean.

A recently proposed model for over-dispersed spike counts in the early visual pathway is the *modulated Poisson* model, in which the stimulus-dependent Poisson spike rate is modulated by a multiplicative stochastic gain variable on each trial (Goris et al.,2014). The model can be written^
1
^:

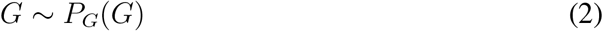

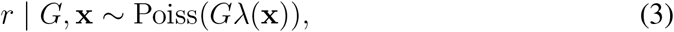

where *P_G_
* is the distribution of a (unobserved) stochastic gain variable *G*, which is assumed to have mean 𝔼[*G*] = 1, and variance var(*G*) = 
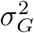
. The conditional distribution of the response given the stimulus requires marginalizing over *G*:

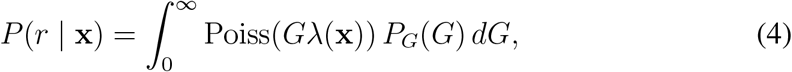

where Poiss(·)denotes the Poisson distribution (Eq. 1). When *G* has a gamma distribution, this integral can be evaluated in closed form and results in a negative binomial distribution, a popular model for over-dispersed spike counts (Pillow & Scott, 2012b; Goris et al., 2014; Linderman et al., 2016). (See Appendix A for a derivation of this relationship).

For any choice of *P_G_
*, the constraint 𝔼[*G*] = 1 ensures that the gain variable has no effect on the marginal mean spike count, meaning that E[*r* [**x**] = *λ*(**x**) as in the pure Poisson model. However, the variance exhibits a quadratic dependence on the spike rate, which can be derived using the law of total variance:

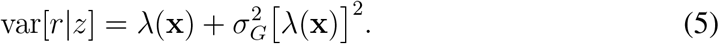

If the gain variable variance 
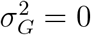
, meaning the gain variable is constant (*G* = 1), the model reverts to Poisson. However, for 
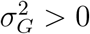
 it exhibits over-dispersion that increases with firing rate, with Fano factor given by

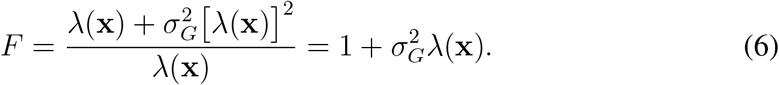

Goris et al. (2014) showed that this model provides a more accurate description of mean-variance relationships than the Poisson model for neurons from multiple visual areas (LGN, V1, V4 and MT). However, as we will show in the following, the quadratic relationship between variance and mean imposed by the Goris model does not accurately apply to all neurons.

## 3 The flexible over-dispersion model

Our flexible over-dispersion model for over-dispersed spike counts consists of a stimulus dependent term *g*(**x**), additive Gaussian noise *n*, and a nonlinear function *f* that transforms (*g*(**x**) + *n*) to a firing rate, followed by Poisson firing. Mathematically it can be written:

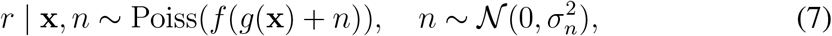

where *f* is a nonnegative, monotonically increasing nonlinearity that ensures nonnegative firing rates. The joint distribution over spike count and latent noise is therefore a product of Poisson and Gaussian distributions,

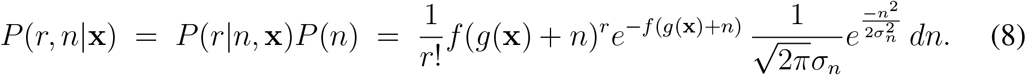

This flexible over-dispersion model can be interpreted in terms of a noisy sub-threshold membrane potential that is transformed by an output nonlinearity to drive Poisson spiking (c.f. Carandini (2004)). In this interpretation, *g*(**x**) is the stimulus tuning curve of the membrane potential, 
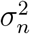
 is the variance of noise affecting in the membrane potential, and *f* (·) is a rectifying output nonlinearity that converts membrane potential to firing rate.

To obtain the probability of a spike count given the stimulus, we marginalize this distribution over the unobserved noise *n*:

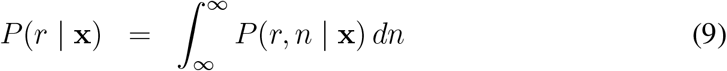

Fig. 1 shows a diagram illustrating the model and several example choices for the non-linearity *f* (·).

The choice of nonlinearity *f* (·) allows the model the flexibility to produce different forms of over-dispersion, that is, mean-variance curves with different shapes. When *f* (·) is exponential, we recover the Goris model framework with multiplicative noise, since exp(*g*(**x**)+*n*) = exp(*g*(**x**))·exp(*n*), meaning the noise multiplicatively modulates with the stimulus-induced Poisson firing rate exp(*g*(**x**)). However, as we will show, other choices of *f* (·) lead to different over-dispersion curves from the quadratic meanvariance curve implied by multiplicative noise, as depicted in Fig. 2.

**Figure 2:**
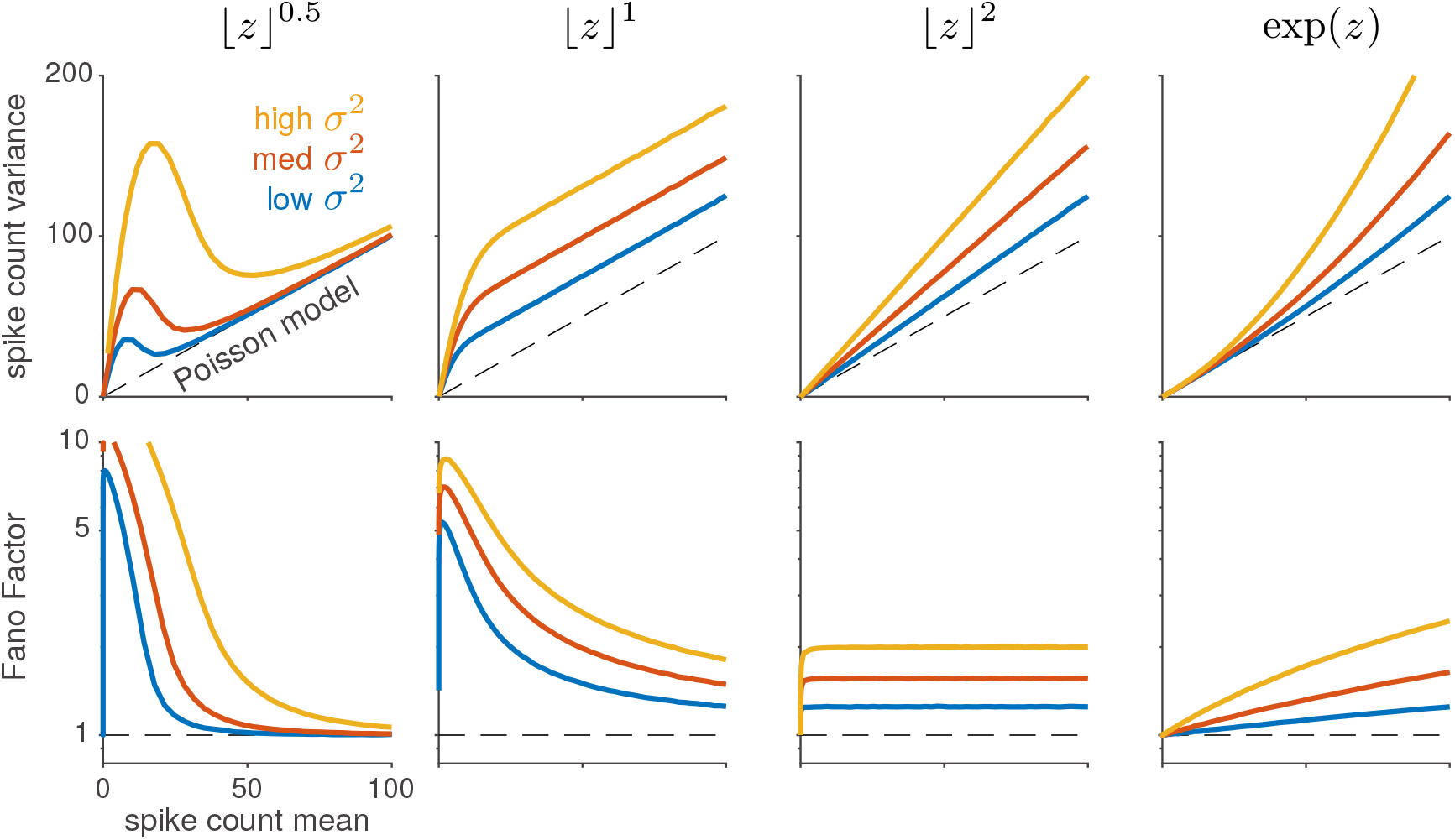
Variance-mean curves for four different choices of nonlinearity *f*, each at three different levels of noise variance 
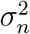
. The top row shows variance as a function of mean, while the bottom row shows Fano factor vs. mean. Note that only the squaring nonlinearity (third column) produces responses of (approximately) constant Fano factor, that is, constant-slope variance-mean curves that pass through the origin. The exponential nonlinearity (fourth column) exhibits variance that grows quadratically with mean, as shown in Goris et al. (2014), and thus has monotonically increasing Fano Factor.

### 3.1 Spike count mean and variance

To characterize the over-dispersion quantitatively, we need to compute the spike count mean and variance as a function of the stimulus drive, or stimulus-induced membrane potential, which we denote *z* = *g*(**x**) for notational convenience. The mean spike count, which we denote *λ*(*z*), can be computed via the law of iterated expectations:

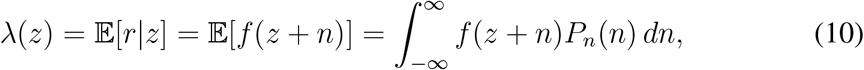

where 
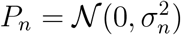
 is the Gaussian distribution of the noise variable *n*.

The spike count variance, denoted 
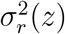
 can be calculated using the law of total variance as

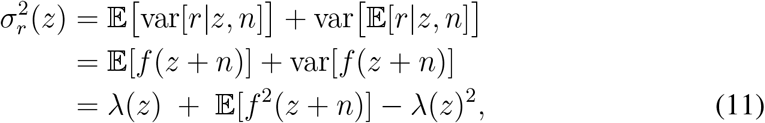

where the expectation in the last line is taken with respect to the Gaussian noise distribution *N* (0, 
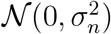
. When 
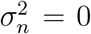
 = 0, the last two terms cancel and the model reverts to Poisson, with variance equal to the mean *λ*(*z*).

For arbitrary choices of *f*, the spike count mean and variance have no closed-form expression and must be computed by numerical integration. However, for several simple choices for *f* we can evaluate these quantities analytically or approximate them closely using the delta method. (See Table. 1).

### 3.2 Capturing different forms of over-dispersion

To illustrate the model’s flexibility, we consider several specific choices for *f*, namely exponential, and rectified power functions, *f* (*z*) = ⌊*z*⌋^
*p*
^ for *p* = {1/2, 1, 2}, where ⌊*z*⌋ = max(*z*, 0). Fig. 2 shows the different mean-variance relationships produced by these four different choices for *f*, at three different values of noise level 
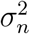
. This demonstrates simple nonlinear transforms that grow at different rates can capture important classes of behavior.

For concave nonlinearities like the rectified square-root, *f* (*z*) = ⌊*z*⌋^0.5^, the spike count variance is actually larger at low spike rates than at high spike rates, and the Fano factor drops rapidly to 1 with increasing spike rate (Fig. 2, column 1). Such responses are thus highly over-dispersed at low spike counts but indistinguishable from Poisson at high spike counts. A linear-rectified nonlinearity, *f* (*z*) = ⌊*z*⌋, on the other hand, produces linear variance-mean curves with unit slope, but with a positive intercept that grows with noise variance 
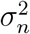
. Although they exhibit a constant degree of over-dispersion relative to Poisson, such neurons have falling Fano factor because the ratio of variance to mean is smaller at high firing rates (Fig. 2, column 2). Interestingly, a rectified squaring nonlinearity, *f* (*z*) = ⌊*z*⌋^2^, gives rise to linear variance-mean curves with slopes that vary as a function of noise variance *σ*
^2^. This is the only model of the four considered that generates constant Fano factor across spike rates (Fig. 2, column 3).

For the exponential nonlinearity, *f* (*z*) = exp(*z*), the spike count mean and variance can both be calculated analytically by exploiting the fact that exp(*n*) has a log-normal distribution (Table. 1). As noted above, this corresponds to the multiplicative noise setting considered by Goris et al. (2014), and exhibits variance that grows quadratically as a function of mean (Fig. 2, column 4). However, we will show in Sec. 5 that this model behaves very differently from the multiplicative model with a gamma-distributed gain variable (i.e., which produces negative-binomial distributed spike counts discussed in Goris et al. (2014)). This indicates that all multiplicative-noise models are not equal; differences in the distribution of the multiplicative noise variable can give rise to major differences in higher-order moments, which in turn can produce very different fits to data.

**Table 1:** Spike count mean and variance for specified nonlinearities. Here Φ(*·*) denotes the normal cumulative density function and 
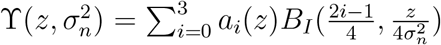
 with *B_I_
* (*·, ·*) denoting the modified Bessel function of the first kind and weighting coefficients *a*
_0_ = *−z*
^2^, 
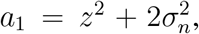
, *a*
_2_ = *z*
^2^, *a*
_3_ = 1. To calculate the approximate variances for *f* (*z*) = *lzJ^p^
* we used the delta method (valid for small 
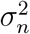
 to obtain a general approximation: 
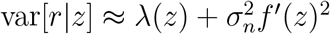

| nonlinearity | mean $\lambda(z)$ | variance $\sigma_r^2$ |
| --- | --- | --- |
| $f(z) = \lfloor z \rfloor^{\frac{1}{2}}$ | $\frac{\sqrt{\pi}}{2\sigma_n\sqrt{-2z}} \exp(-\frac{z^2}{2\sigma_n^2}) \Upsilon(z, \sigma_n^2)$ | $\approx \lambda(z) + \sigma_n^2 \frac{1}{\lambda(z)^2}$ |
| $f(z) = \lfloor z \rfloor$ | $\frac{\sigma_n}{\sqrt{2\pi}} \exp(-\frac{z^2}{2\sigma_n^2}) + z\Phi\left(\frac{z}{\sigma_n}\right)$ | $\approx \lambda(z) + \sigma_n^2$ |
| $f(z) = \lfloor z \rfloor^2$ | $\frac{\sigma_n z}{\sqrt{2\pi}} \exp(-\frac{z^2}{2\sigma_n^2}) + (z^2 + \sigma_n^2)\Phi\left(\frac{z}{\sigma_n}\right)$ | $\approx (1 + 4\sigma_n^2)f(z)$ |
| $f(z) = \exp(z)$ | $\exp(z + \frac{1}{2}\sigma_n^2)$ | $\lambda(z) + (\exp(\sigma_n^2) - 1)\lambda(z)^2$ |

It is also worth noting that a model with latent Gaussian noise and exponential nonlinearity is also the default model considered in an extensive literature on factor or latent linear dynamical system models for multivariate spike count data (Macke et al., 2011; Buesing et al., 2012; Archer et al., 2014; Rabinowitz et al., 2015; Ecker et al., 2016; Gao et al., 2016; Zhao & Park, 2017). The majority of this literature, however, has focused on capturing covariance across time or across neurons, and has devoted less attention to the issue of how count variance grows with mean for single neurons.

While solutions given in Table 1 and Fig. 2 are useful for building intuition, in practice it makes sense to choose a parametrized function whose parameters can be adapted to capture a variety of different behaviors. For the general case *f* (*z*) = ⌊*z*⌋^
*p*
^, *p >* 0, *p ≠* 1 we can use the delta method to derive the approximate variance as

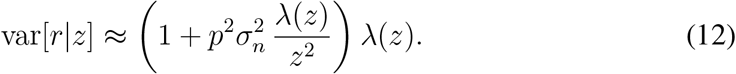

Note that when 
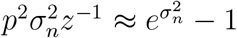
, this expression for the variance mimics the mean-variance relation achieved by the exponential nonlinearity (Table 1). Thus, for larger powers of *p >* 2, the rectification nonlinearity can reasonable approximate super-linear mean-variance relationships over the range of means *λ*(*z*) such that 
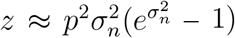

^−1^.

Additionally, as optimizing these parameters to fit neuron firing counts will require calculating various derivatives, a smooth function is also beneficial for stability in the fitting process. We find that a useful alternative to the rectified power function given above is the soft-rectification power function, i.e. a soft rectification raised to a power *p*:

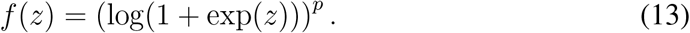

By fitting both *p* and the unknown latent variance 
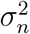
, we find that this model is rich enough to capture a wide variety of over-dispersion behaviors.

Lastly, we note that despite *λ*(*z*) and 
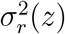
 often requiring numerical integration to obtain, we can guarantee certain properties on the parametric curve, as parametrized by *z*. First, we can guarantee that both *λ*(*z*) and 
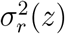
 are smooth functions of *z* (although the derivatives may be discontinuous). Second, so long as the function *f* (·) is positive and monotonically increasing, *λ*(*z*) is also a monotonically increasing function of *z*. This indicates that 
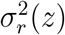
 can truly be considered a function of *λ*(*z*) since each value of *λ*(*z*) maps onto only one value of 
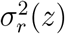
. No similar guarantee could be made for 
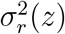
, however, indicating that local minima can exist in the mean-variance relation. Local minima occur when the function *f* (·) has regions that are concave and have small derivatives, as occurs for *f* (*z*) = ⌊*z*⌋ *
^p^
* (see Fig. 2). In fact, if *f* (·) were constructed to have multiple concave regions with small derivatives, multiple local minima and maxima could be present in the mean-variance relationship. Finally, we note that since 𝔼[*f* ^2^(*z* + *n*)] *≥* 𝔼[*f* (*z* + *n*)]^2^, we always have 
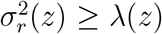
 *≥ λ*(*z*), indicating that while the amount of over-dispersion may vary significantly, out model always produces greaterthan-Poisson spike count variance.

## 4 Model fitting

In order to apply the flexible over-dispersion model to neural data, we need a tractable method for fitting the model parameters. Consider a dataset composed of a set of stimuli *X* = {**x**
_
*i*
_} and the measured spike counts of a single neuron *R* = {*r_ij_
*} where *r_ij_
* denotes the *j*’th response of the neuron to stimulus **x**
_
*i*
_, for *i∈*{1, …, *N*}. The model parameters *θ* consist of a stimulus-dependent term for each stimulus, denoted *z_i_
* = *g*(**x**
_
*i*
_), the noise variance 
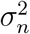
, and (optionally) the exponent *p* governing the soft-rectification power nonlinearity. (We will consider the variable *p* to stand for any parameters governing the nonlinearity, although for the exponential nonlinearity there are no additional parameters to fit).

The likelihood is given by a product of conditionally independent terms, one for each spike count:

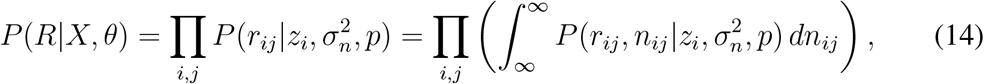

where *θ* = {{*z_i_
*},*σ*
^2^, *p*} is the full set of model parameters, and 
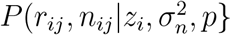
 is the joint distribution of the spike count *r_ij_
* and latent noise *n_ij_
* for the *j*’th response to stimulis **x**
_
*i*
_, which is given by a product of Poisson and Gaussian distributions (eq. 9).

Because the integral over the latent noise *n* is intractable, we compute the likelihood using the Laplace approximation. This method uses a surrogate Gaussian distribution to approximate the posterior of *n* given *r* around its mode:

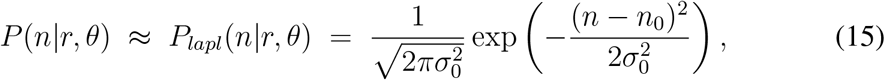

where mean *n*
_0_ is the mode of *P* (*n|r, θ*), which we find via a numerical optimization of log *P* (*r, n|θ*) for *n*, and variance 
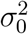
 is the inverse of the Hessian (2nd derivative) of the negative posterior log-likelihood,

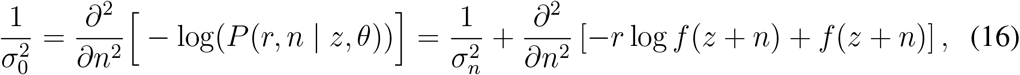

evaluated at mode *n* = *n*
_0_. If *f* is exponential, Equation (16) simplifies to 
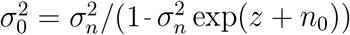
.

To evaluate the likelihood, we can replace the true posterior with the Laplace approximation to obtain a tractable form for the joint distribution, *P* (*r, n | θ*) *≈ P* (*r | θ*) *· P_lapl_
*(*n | r, θ*). We can then solve this expression for the desired likelihood, yielding:

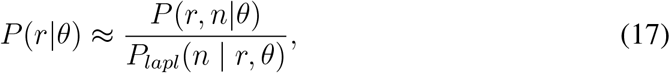

where the numerator is the exact joint probability of *r* and *n* given *θ*, and the denominator is the approximation to the posterior over *n* given *r*. Evaluating the right-hand-side at *n* = *n*
_0_ (where the Laplace approximation is most accurate) gives the following expression for the likelihood:

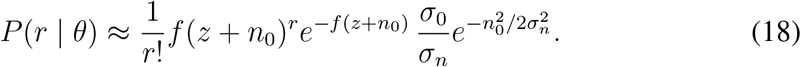

Note that the parameters of the Laplace approximation, *n*
_0_ and 
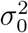
, are functions of *θ*, since they depend on numerical optimization of *P* (*n, r|θ*) for *n* at a given *θ*. Thus, maximum likelihood parameter estimation, which involves maximizing the log of Equation (14) for *θ*, requires we update these per-response parameters defining the Gaussian distribution for each noise term *n_ij_
* given *r_ij_
* and *θ* every time we change *θ*.

To demonstrate that this method yields an accurate approximation to the log-likelihood, we compare the true computed log-likelihood (via numerical integration) with the Laplace approximation. Fig. 3 depicts the log-likelihood landscapes for different spike counts (*r∈*{0, 2, 10, 50}) for both the numerically integrated log-likelihood and the Laplace approximation. The ratio between numerically computed and Laplace based log-likelihoods (rightmost column) demonstrates that errors are greatest at small spike counts (where the Poisson likelihood deviates most severely from Gaussian), but is negligible even in this regime for the exponential nonlinearity.

**Figure 3:**
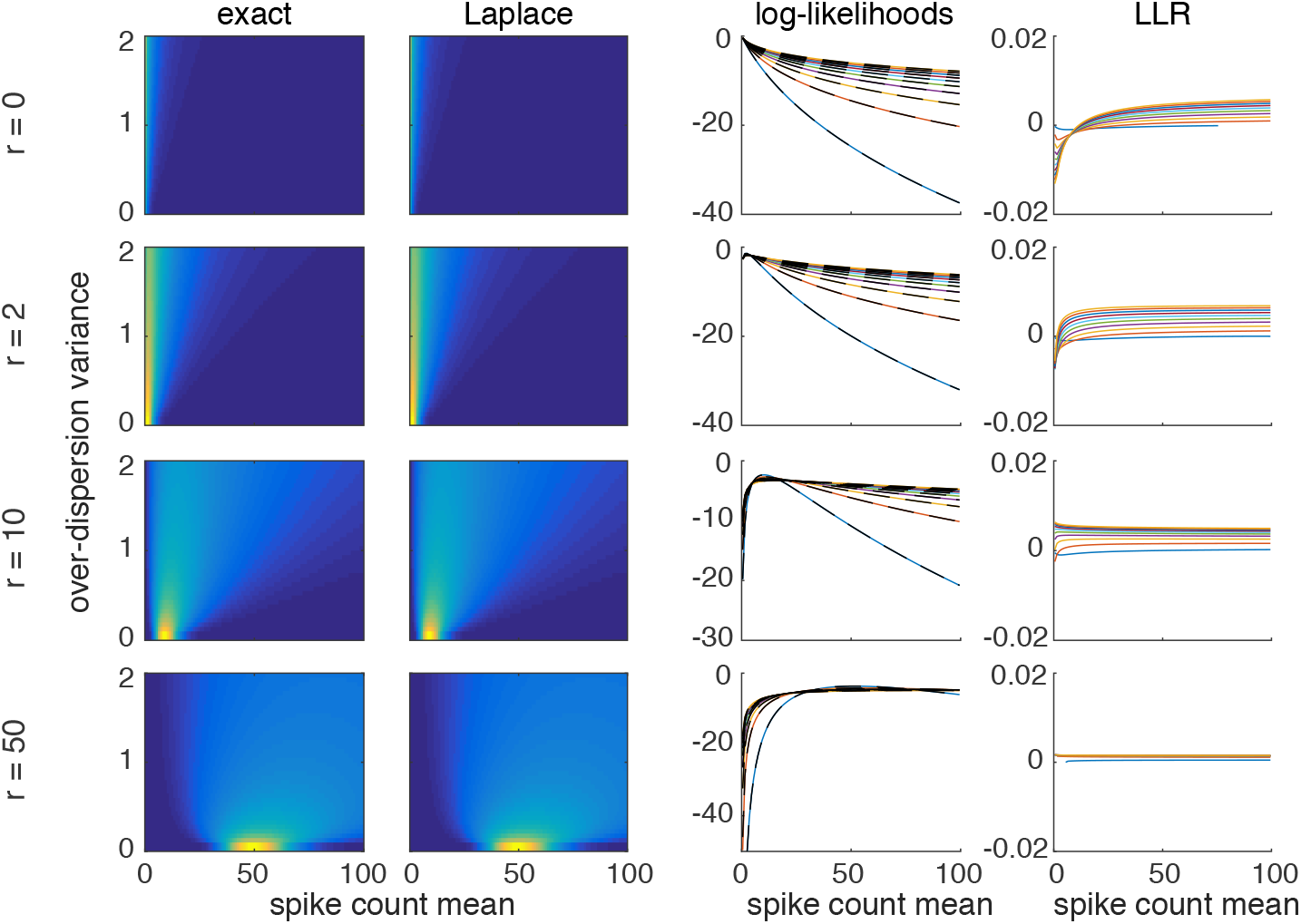
Comparison of exact and Laplace approximation-based evaluation of model likelihood function under exponential nonlinearity. Left two columns: image plots show log-likelihood function log *P(r|z*, 
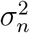
 for spike counts *r* = 0, 2, 10, and 50, as a function of transformed parameters 
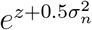
 (mean spike count) and (
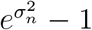
 *−* 1) (over-dispersion variance for Goris model). The third column shows horizontal slices through the two surfaces at different over-dispersion levels (colored traces for exact; black dashed traces for Laplace), showing the two surfaces are virtually indistinguishable on the scale of the log-likelihood values. The last column shows the ratio between the true (numerically computed) and Laplace-based log-likelihood, showing there is less than a 0.01% error in the approximation across the entire range of parameter settings at these spike counts. We note that the approximation error is greatest at low spike counts, where the Poisson log-likelihood is most non-Gaussian.

We can use the above methodology to now address the particular case where *f* (·) is the soft-rectification raised to a power 
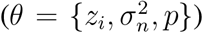
. With this nonlinearity, we obtain the update equations (see also Appendix B):

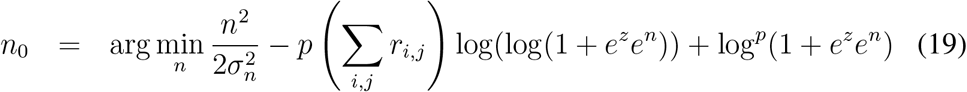

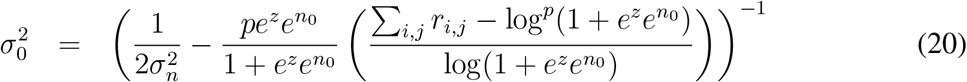

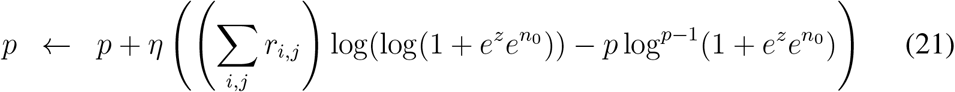

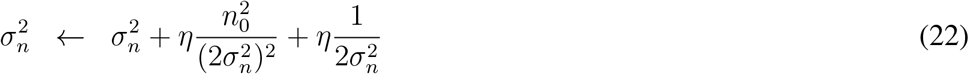

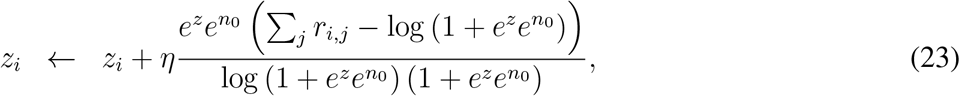

where *η* is the gradient descent step-size. For the case where *f* (·) is an exponential function, these expressions reduce further, and the full model fitting method is provided in Appendix B.

## 5 Results: V1 spike counts

To test the ability of the flexible over-dispersion model to capture the variability of real neurons, we fit to data collected from macaque primary visual cortex V1 (Graf et al., 2011) (one of the datasets considered in Goris et al. (2014)). This dataset contained five recording sessions with multi-electrode array recordings of neural population responses to 72 distinct visual grating orientations. A total of 654 neurons were recorded, and each orientation was presented 50 times. Trials consisted of drifting grating stimulus presentations for 1280ms, with a 1280ms rest period in between. From this dataset, we chose the 112 well-tuned neurons for model fitting. Well-tuned neurons were determined as in previous studies by applying a threshold to the difference between the minimum and maximum mean firing rate over all stimuli orientations (Graf et al., 2011).

We fit the model parameters using the Laplace-approximation based method described in Section 4, for both the soft-rectification-power function (“soft-rect-p”, eq. (13)) and the exponential nonlinearity. For comparison, we also fit the data with the negative-binomial (NB) model from (Goris et al., 2014). All model fits had 72 parameters {*z_i_
*}t o describe the stimulus drive for each orientation and a noise variance parameter 
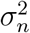
, with the soft-rect-p model requiring an extra exponent parameter *p*. Example mean-variance curves (as well as example draws from the model with the inferred parameters) are plotted against the true data in Fig. 4. While for some neurons the negative binomial model and our model resulted in near-identical mean-variance curves, for other neurons the flexibility of the soft-rectification function produced better qualitative fits. Fig. 5 depicts the range of mean-variance relationships inferred for a sampling of neurons in the data set. The range of curves observed for even this single data set implies that a single mean-variance relationship is insufficient to capture the entire range of neural variability. Fig. 5 further emphasizes this point by showing histograms of the inferred parameters (in the case of soft rectification this includes the latent noise standard deviation *σ_n_
* and the soft-rectification power *p*).

**Figure 4:**
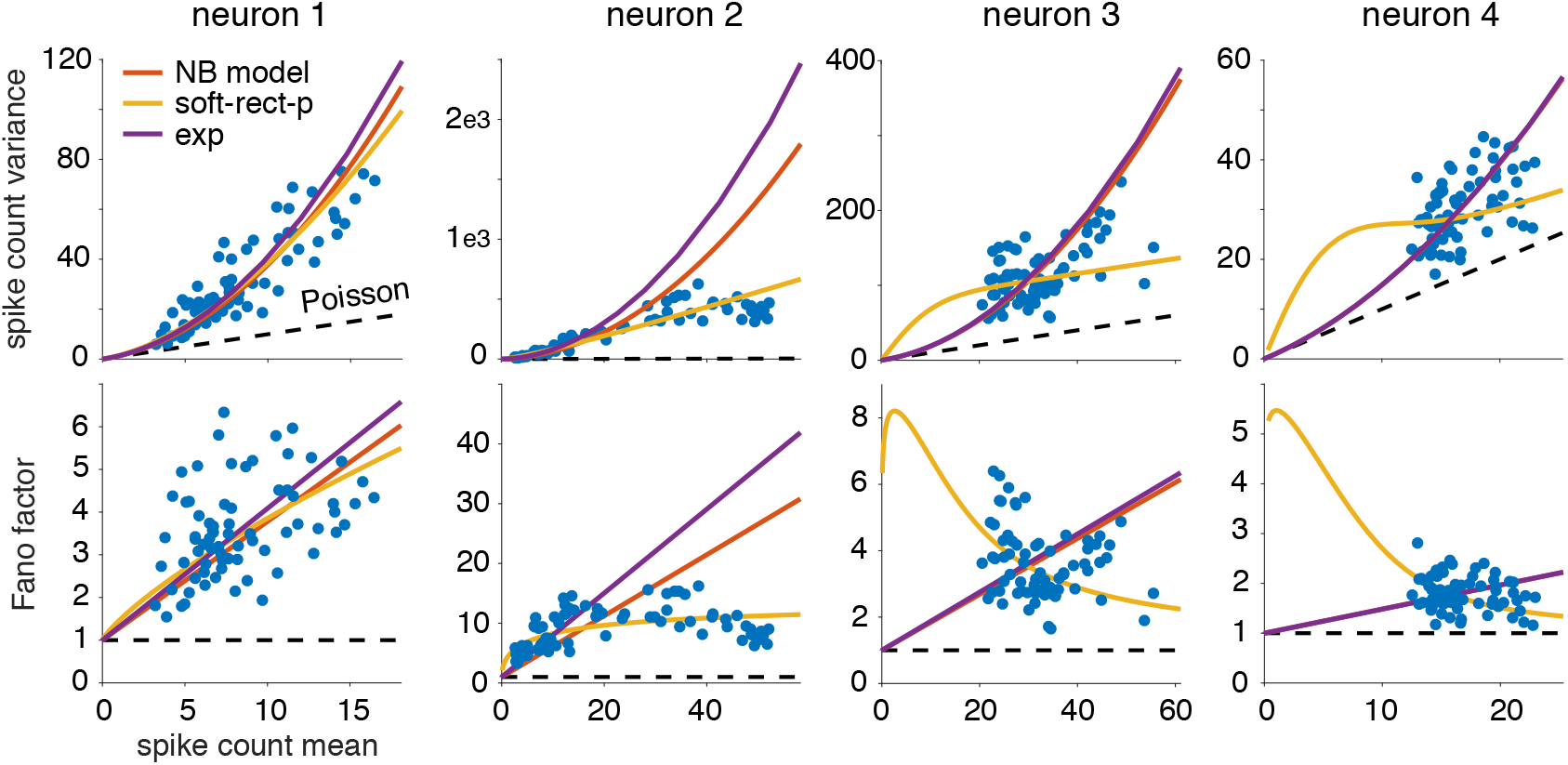
Example fits to V1 neurons. Each column shows data from a single neuron, with upper plot showing variance vs. mean, and lower plot showing Fano factor vs. mean. Empirical spike count mean and variances (blue dots) were computed for over 50 repetitions for each of 72 different oriented grating stimuli (Graf et al., 2011). For each neuron, we fit three different models by maximum likelihood using all 50 × 72 = 3600 spike count responses. The negative binomial model (red line) and flexible over-dispersion model with exponential nonlinearity (yellow) both had 73 parameters (an input level for each of the 72 orientations, and the noise variance 
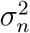
), while the flexible over-dispersion model with soft-rectification power nonlinearity (yellow) had 74 parameters (including power parameter *p*). For all four neurons shown, the flexible over-disperion model with soft-rectification power (soft-rect-p) nonlinearity achieved the best model fit as measured by the Akaike information criterion (AIC). The different distributions of the data in each figure shows that there is significant diversity in neural spiking, even within V1. When the mean-variance data behavior is decidedly not quadratic (neurons 2,3, and 4), the soft-rect-p model can adapt to the wide range of statistics. Interestingly, when the data behavior does look quadratic (neuron 1), the soft-rect-p automatically recovers the quadratic behavior modeled by the NB and Exp models. While in this case the soft-rect-p model still had the best AIC score, the difference from the NB AIC value for this example is only 3, indicating that both models approximately recover the same behavior.

**Figure 5:**
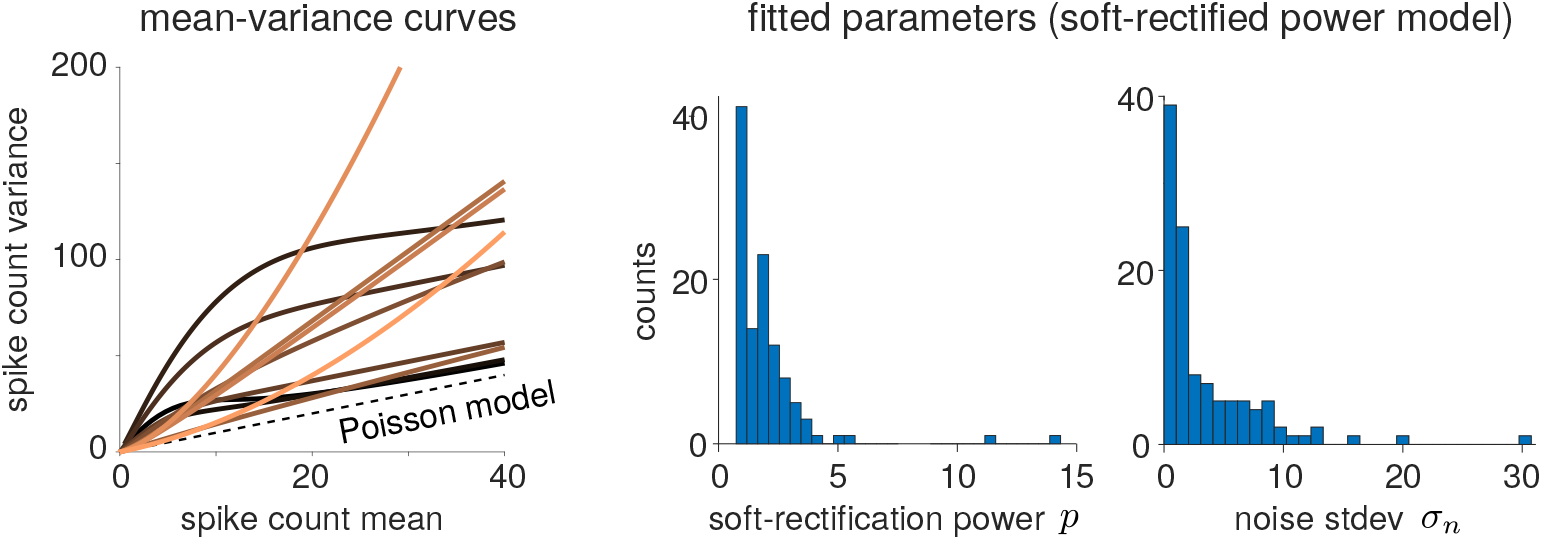
Fits of hierarchical over-dispersion model with soft-rectified power nonlinearity. Left: Selected variance-mean curves obtained from fits to individual neurons in V1 (each trace represents the mean-variance relationship for a single neuron). Curves are colored in order of increasing exponent power *p* (lighter lines have higher *p*). Center: Histogram of the distribution of inferred soft-rectification power parameters *p* for neurons in V1 demonstrates the required flexibility in fitting over-dispersion behavior. Right: the distribution in inferred latent noise standard deviations *σ_n_
* shows the range in the amount of over-dispersion noise in the population of V1 neurons.

We quantified the performance of different models by comparing the Akaike Information Criterion (AIC) across all three fits (power soft-rectification, exponential, and negative binomial). The calculated AIC values, displayed as histograms in Fig. 6, demonstrate that the softrect-p model provided the best fit for the majority of neurons, achieving the highest AIC values for ≈83.4% of the 112 neurons analyzed. This result indicates that despite the AIC penalization for requiring an additional parameter, the softrect-p fit best modeled approximately ≈ 83% of the tuned V1 neurons in the Graf data set. Interestingly, we also note that the exponential nonlinearity model, a model commonly used as an inverse link function in machine learning and neuroscience (Nelder & Baker, 1972) was a *worse* model for 65.8% of the neurons according to AIC, even having performed worse than the NB model (as shown in Fig. 6). This result may seem contrary to the fact that the model with exponential nonlinearity belongs to the same class of multiplicative-noise models as the negative binomial and thus also exhibits a quadratic mean-variance curve. Fig. 4 demonstrates this behavior, showing that data fits to both quadratic models can still differ significantly in their curvature. The implication here is that while the full mean-variance curve may provide basic over-dispersion properties, it is insufficient to describe a neural spiking model. Specifically, model fitting with a full probabilistic model implies assumptions on higher-order moments that will effect the mean-variance curve fit.

**Figure 6:**
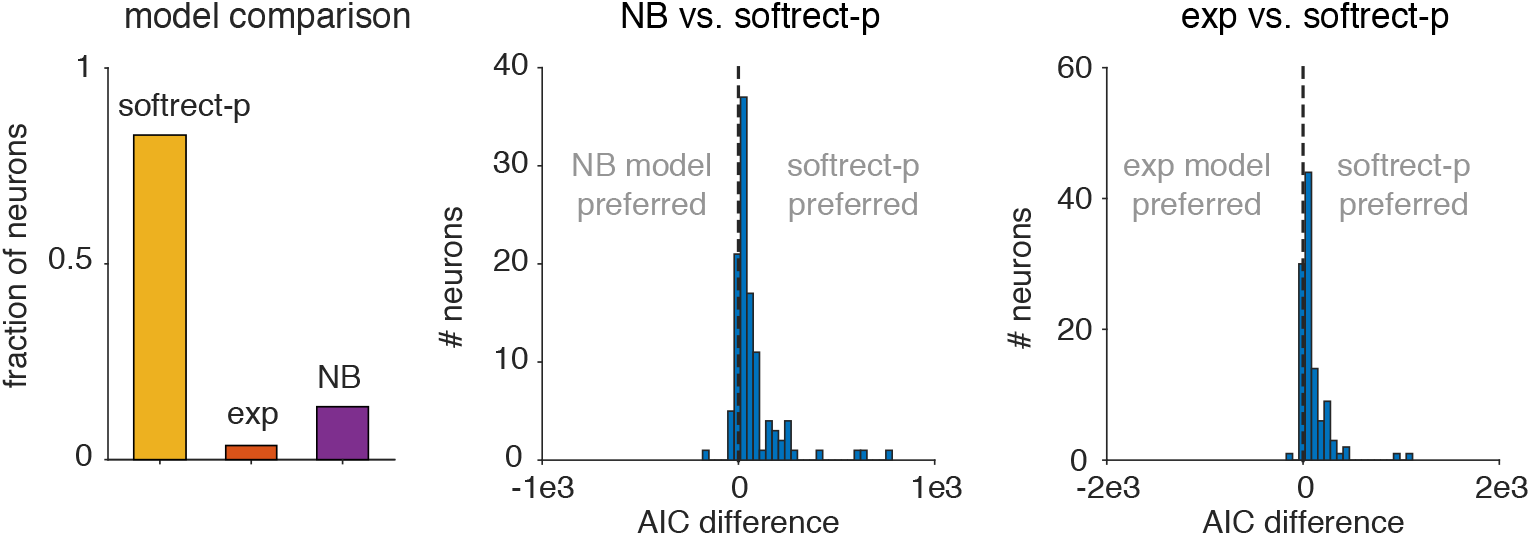
Model comparison between negative binomial (NB), and flexible overdispersion model with exponential and soft-rectification power nonlinearity using AIC. The soft-rectification power (with fitted power *p*) performed best for 82% of the neurons, while the NB model performed best for 13.5% (left panel). Pairwise comparisons (middle and right panels) show the distribution of AIC differences across all neurons for pairs of models, in agreement with the overall comparison results.

To emphasize the potential insufficiency of the mean-variance curve as a model summary, Fig. 7 shows an example case where the exponential nonlinearity yielded a very different mean-variance curve than the negative binomial model, despite the fact that both are multiplicative noise models defined by a stochastic gain variable with a mean of 1. Specifically, for this neuron, both the exponential and NB models seem to have significantly over-estimated the variance as a function of the mean. The exponential fit over-estimation was so extreme that the curve for the exponential fit is barely visible in the mean-variance plot. After ruling out potential numerical or approximation error with the Laplace approximation technique, we observed that while the mean-variance curves did not match the data, the full count-distributions did seem to match the data. The upper-right and upper-left plots show that overall both models fit the data distributions for both high and low mean spike-counts. Plotting the spike-count distributions on a log-scale, however, shows that despite matching the data well at low firing rates, both models have higher tails than what would be expected. These heavy tails contributed directly to the over-fitting of the spike-count variance. In fact, the exponential nonlinearity’s spike-count distribution has a much heavier tail than even the NB model, correlating with how the exponential nonlinearity model nonlinearity modeled the variance to a higher degree. To understand why the erroneous fits were obtained, we observe that up to a certain value of spike counts, the tails of the data distribution do actually look heavy-tailed. In fact the log-distributions shown in the lower-left and lower-right panels of Fig. 7 do follow the data. Thus heavy tails stemmed from matching the data in the ML estimation. Unlike the data, however, our models are unbounded, indicating that heavy tails could impact the model variance calculations in ways that would not affect the data fits. This analysis demonstrates that attempting to fit the higher-order moment of the model (as ML-type methods do) may be detrimental to accurately fitting the mean-variance curve.

**Figure 7:**
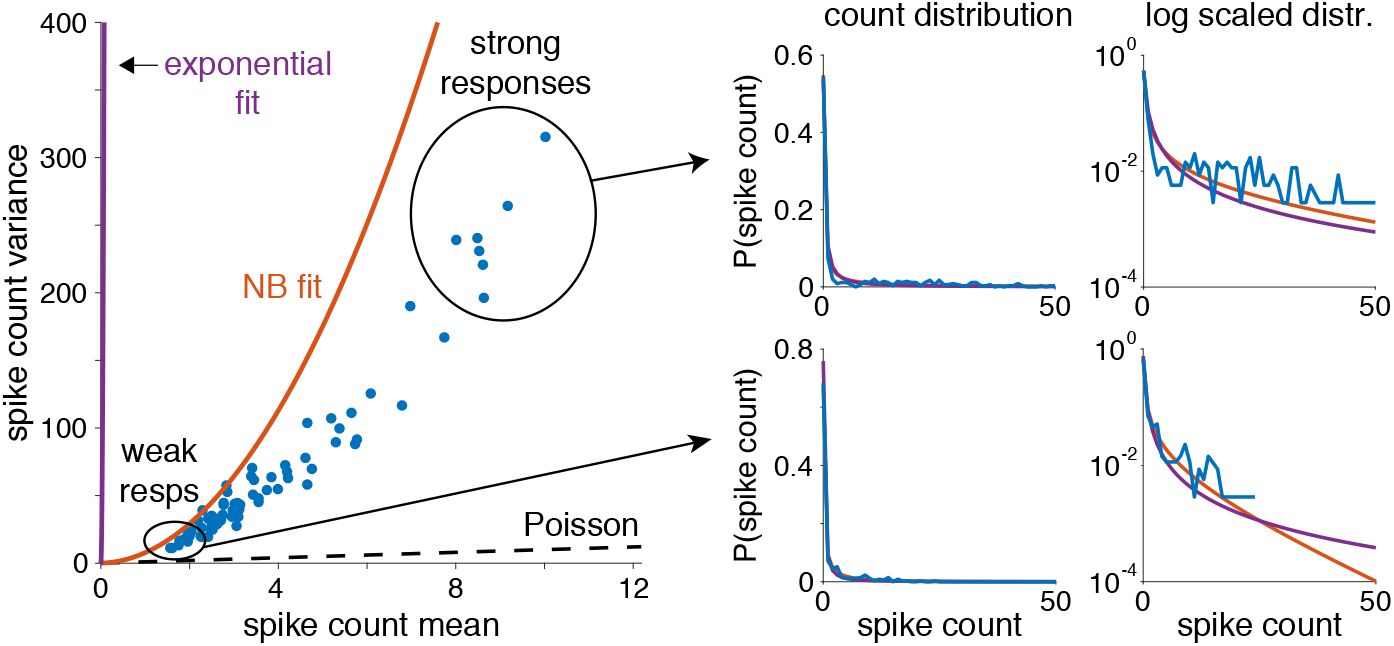
Heavy tails can cause significant model mismatch when fitting over-dispersed models. An example neuron where model fits (left) produced significant overestimation of the variance. The extreme case here produced an NB fit (red curve) with higher variance and an exponential fit (purple curve) that barely reaches the data points (blue dots indicate mean and variance over the 50 trials per stimulus) in the mean-variance plot. Both fits, however, do seem to match the basic statistics of the data when viewed as a distribution. Specifically, when restricted to the stimuli trials eliciting either low or high levels of activity (right plots; bottom plots for low activity and top pots for high activity) we see that the fits did match the spike count distributions (linear y-axis scale on the left and logarithmic scale on the right for clarity) over the support where data was available. The tails of the model distributions, however, differed significantly in both cases, indicating that the model fits are imputing the distribution of spike counts in for unobserved counts. Thus the inherent bias of the models manifested as the observed model mismatch.

## 6 Application: adaptive closed-loop experiments

To illustrate one of the potential uses of the flexible over-dispersion model, we developed an application to adaptive stimulus selection for closed-loop neurophysiology experiments. Such methods, known as *active learning* in machine learning or *adaptive optimal experimental design* in statistics, seek to select stimuli based on the stimuli and responses observed so far during an experiment in order to characterize the neuron as quickly and efficiently as possible (Lindley, 1956; Bernardo, 1979; MacKay, 1992; Chaloner & Verdinelli, 1995; Cohn et al., 1996; Paninski, 2005). Adaptive stimulus selection is particularly useful in settings where data are limited or expensive to collect, and can substantially reduce in the number of trials needed for fitting an accurate model of neural responses (Paninski et al., 2007; Benda et al., 2007; Lewi et al., 2009; DiMattina & Zhang, 2011, 2013; Bölinger & Gollisch, 2012; Park & Pillow, 2012; Park et al., 2014; Pillow & Park, 2016).

Here we introduce a method for adaptive stimulus selection for estimating a neuron’s multi-dimensional tuning curve or “firing rate map” under the flexible over-dispersion model. Our method involves computing the posterior distribution over the tuning curve given the previously observed responses in an experiment, and selects the stimulus for which the value of the tuning curve has maximal posterior variance. We illustrate the performance gain using this adaptive learning method for estimating color tuning curves of V1 neurons recorded in awake, fixating monkeys (Park et al., 2014). For simplicity, we use the exponential nonlinearity *f* (*z*) = exp(*z*), which has the advantage of having analytic expressions for the mean and variance of the spike count given an input level *z* and noise variance 
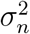
,

### 6.1 GP tuning curve model with flexible over-dispersion

Building on prior work (Rad & Paninski, 2010; Park et al., 2014; Pillow & Park, 2016), we model tuning curves by placing a GP prior over the input function *g*(**x**). Here *g*(**x**) can be considered log of the tuning curve plus a constant, since the tuning curve (expected spike count given a stimulus) for the exponential nonlinearity is 
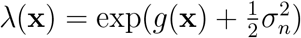
 (Table 1).

The GP specifies a multivariate normal distribution over the input function values {*g*(**x**
_1_), …, *g*(**x**
_
*n*
_)} at any finite collection of stimulus values *{*
**x**
_1_, *…* **x**
_
*n*
_}, namely:

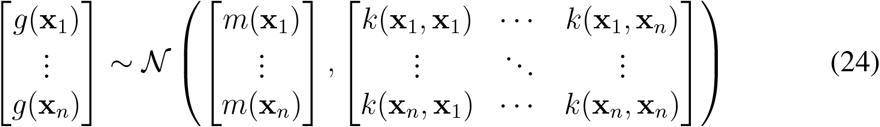

where *m*(**x**) is the *mean function*, specifying the *a priori* mean of the function values, and *k*(**x**, **x**
^
*/*
^) is the *covariance function*, specifying the prior covariance for any pair of function values *g*(**x**), *g*(**x**
^
*/*
^). Here we use a zero mean function, *m*(**x**) = 0, and a Gaussian or *radial basis function* (RBF) covariance function

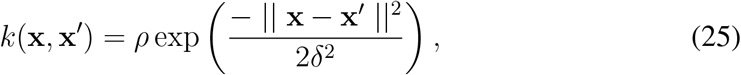

which is controlled by two hyperparameters: the marginal prior variance *ρ* and length scale *δ*. Increasing *ρ* increases the expected prior range of tuning curve values (i.e., increasing the spread between minimal to maximal log firing rates), while increasing *δ* increases its degree of smoothness. Functions *g*(·) sampled from a GP with this covariance function are continuous and infinitely differentiable.

We combine the GP prior with the likelihood from the flexible over-disperion model (eq. 9) to obtain a complete model for Bayesian adaptive stimulus selection. The complete tuning curve model can therefore be summarized:

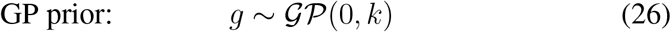

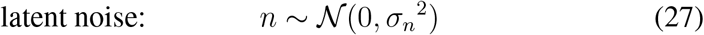

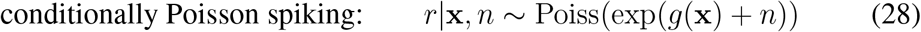

The necessary steps in our stimulus selection algorithm include include updating the posterior over *g*(·) after each trial, updating hyperparameters using the Laplace approximation based marginal likelihood, and selecting the stimulus for which the tuning curve has maximal posterior variance.

### 6.2 Inference and Laplace approximation

The tuning curve inference problem here is similar to that of fitting the flexible overdispersion model (Sec. 4), except that we have added a prior over the values of *g*(**x**) that encourages smoothness and shrinks them toward zero. After each trial, we maximize marginal likelihood under the Laplace approximation to update the hyperparameters (*ρ, δ*) governing the GP prior’s scale and smoothness, as well as the noise variance 
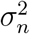
.

Let 𝒟_
*t*
_ = {**x**
_1_, *r*
_1_), *…*, (**x**
_
*t*
_, r_
*t*
_)} denote the dataset after *t* trials, which is simply a paired list of stimuli and spike counts collected so far in the experiment. For the purposes of our derivation, let **z** = [*g*(**x**
_1_), *…, g*(**x**
_
*t*
_)] denote the stimulus input values for the observed stimulus set **X** = [**x**
_1_, *…*, **x**
_
*t*
_], and let **n** = [*n*
_1_, *…, n_t_
*] and **r** = [*r*
_1_, *…, r_t_
*] be the unobserved Gaussian input noise and observed spike counts on each trial, respectively.

The first step in the inference procedure is to maximize the log of the joint posterior over **z** and **n**, also known as the *total-data posterior*, after each trial:

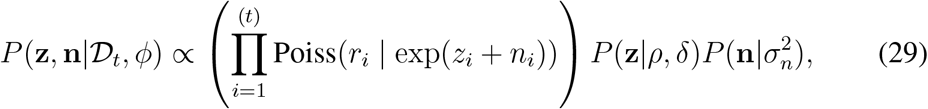

where *P* (**z |**
*ρ, δ*) = (0, *K*) is the GP-induced Gaussian prior over the function values (eq. 24) with covariance matrix *K_ij_
* = *k*(**x**
_
*i*
_, **x**
_
*j*
_), and 
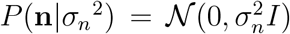
 is the distribution of the (unobserved) over-dispersion noise for each trial. The full set of hyperparameters is denoted 
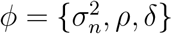

This optimization is simplified by considering a change of variables to take advantage of the fact that the Poisson likelihood relies only on the sums of *z_i_
* + *n_i_
* for each trial. Therefore, let us define:

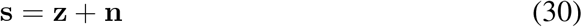

so that **s** represents the total input to *f* (·) on each trial, and the latent noise is now **n** = **s** – **z**.

The total-data log posterior can now be written in terms of **s** and **z**:

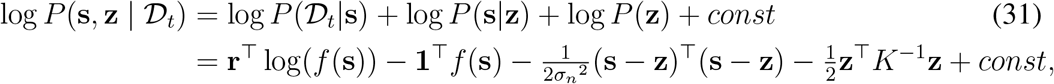

where **1** is a length-*t* vector of ones, and *const* denotes constants that are independent of **s** and **z**. Note that the total-data log posterior is quadratic in **z**, meaning that the conditional maximum in **z** given **s** can be obtained analytically:

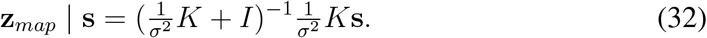

We can substitute this expression for **z** in (eq. 31) to obtain the profile likelihood:

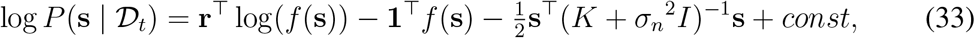

which we can optimize efficiently to obtain the MAP estimate:

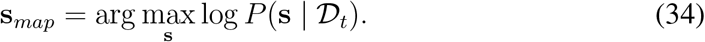

Note the vector **s** grows by one element after every trial, but this optimization can be made efficient by initializing the first *t −* 1 elements of the vector to its value from the previous trial. The joint optimum of the total data log-posterior can be obtained by plugging **s**
_
*map*
_ into (eq. 32) to obtain **z**
_
*map*
_, and setting **n**
_
*map*
_ = **s**
_
*map*
_ – **z**
_
*map*
_.

The Laplace approximation of the total-data posterior can then be computed as:

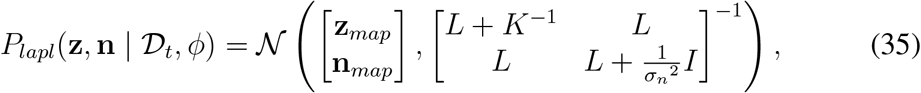

where the blocks of the inverse covariance come from the negative Hessians of the total-data log posterior:

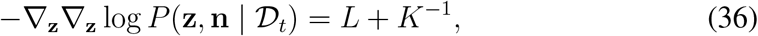

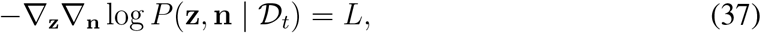

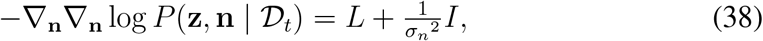

where *L* = – **∇_z_∇ _z_
** log *P* (D_
*t*
_ **z**,*|* **n**) = diag[exp(**z** + **n**)] is a diagonal matrix with second derivatives from the Poisson term, all evaluated at **z** = **z**
_
*map*
_ and **n** = **n**
_
*map*
_.

From the joint Gaussian posterior (eq. 35), we can compute the marginal posterior over **z**, the tuning curve inputs at the stimuli in our training set:

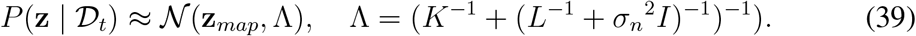

For stimuli not in our training set, denoted **x**
^
*∗*
^, (e.g., candidate stimuli for the next trial), we can compute the marginal posterior distribution over the input value *z^∗^
* = *g*(**x**
^
*∗*
^) using the Gaussian identities that arise in GP regression (Rasmussen & Williams, 2006):

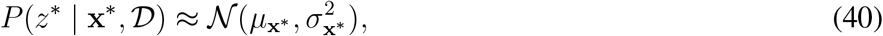

where

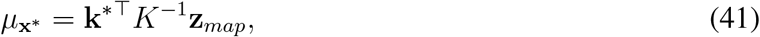

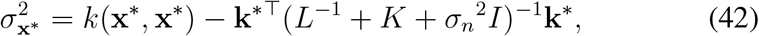

where 
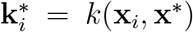
 is the GP covariance function evaluated at the *i*’th presented stimulus **x**
_
*i*
_ and test stimulus **x**
^
*∗*
^.

### 6.3 Hyperparameter optimization

To optimize hyperparameters 
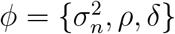
 governing the GP prior and the overdisperion noise, we use the Laplace approximation to compute the marginal likelihood:

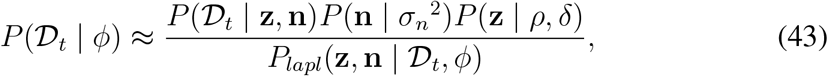

evaluated at **z** = **z**
_
*map*
_ and **n** = **n**
_
*map*
_, where the denominator is the Gaussian approximation to the posterior (eq. 35). This gives:

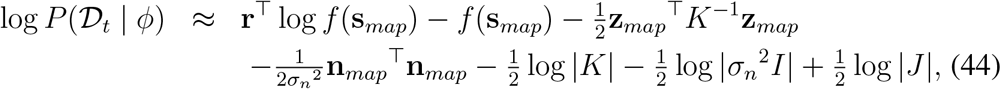

where **s**
_
*map*
_ = **z**
_
*map*
_ +**n**
_
*map*
_, and *J* is the covariance matrix of the Laplace approximation in (35), with determinant given by 
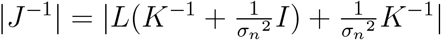
.

Fig. 8 shows the consequences of over-dispersed spike responses on the estimation of tuning curves and their hyperparameters. We generated a 2D map firing rate map as a mixture of Gaussian bumps (left), and simulated spike counts from the flexible over-dispersion model (with exponential nonlinearity) at a random collection of 2D locations. We then performed inference using either the Poisson model (center) or flexible over-dispersion model (right). We found that the Poisson estimate suffered from systematic under-estimation of the GP length scale, resulting in an estimate that is significantly rougher than the true map. In essence, the Poisson model attributes super-Poisson variability of the responses as reflecting structure in the firing rate map itself rather than noise, thus necessitating a shorter length scale *δ*.

**Figure 8:**
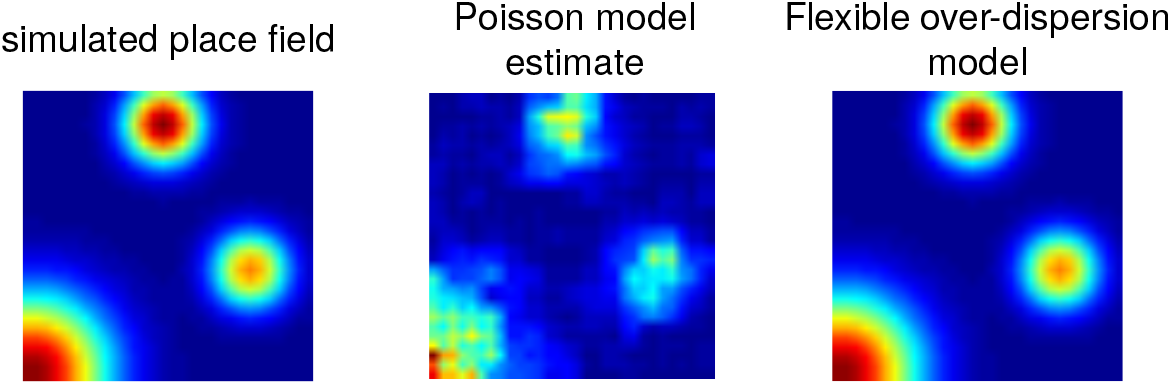
Proof-of-concept experiments on simulated data show that accounting for super-Poisson firing statistics can increase accuracy in recovering unknown firing maps. The true simulated firing map (left plot) drove a model with over-dispersed spiking. We used the simulated spike counts to infer the underlying firing rate map using a Poisson response model (center plot) and the flexible over-dispersion model (right plot). The center plot demonstrates how the Poisson assumption can impair estimates of tuning curve smoothness; accounting for over-dispersed spike count statistics improves the estimate substantially by allowing for more accurate estimation of the GP length scale hyperparameter *δ* that governs smoothness.

### 6.4 Adaptive stimulus selection method

To adaptively select stimuli during an experiment, we search for the stimulus for which the posterior variance of the tuning curve is maximal. This approach, commonly known as uncertainty sampling, is motivated by the idea that we would like to minimize posterior uncertainty about the tuning curve at all points within the stimulus range.

To recap, the tuning curve given given the input function *g*(·) is the expected spike count given a stimulus:

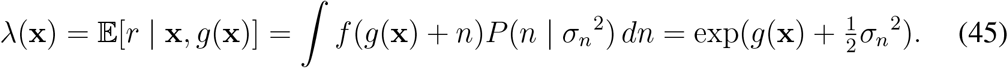

However, during inference from data we have uncertainty about *g*(·), which is characterized by the (approximate) Gaussian posterior derived in the previous section (eq. 40). To compute the posterior mean and variance over the tuning curve *λ*(**x**
^
*∗*
^) at any candidate stimulus **x**
^
*∗*
^, we have to transform the posterior distribution over *g*(**x**
^
*∗*
^) through the nonlinearity *f*.

Letting *z^∗^
* = *g*(**x**
^
*∗*
^) as before, the mean and variance of the tuning curve are given by:

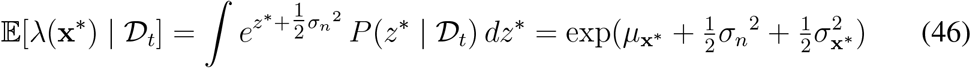

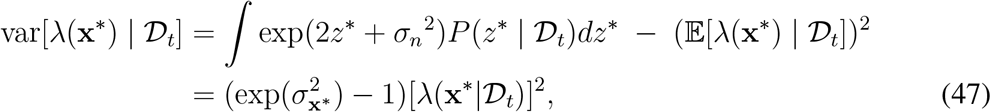

with stimulus-specific mean and variance *µ*
_
**x**
_
*∗* and 
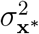
 given in (eqs. 41, 42). In practice, our method involves computing posterior variance (eq. 47) for a grid of candidate stimuli {**x**
^
*\**
^} and selecting the one for which posterior variance is largest:

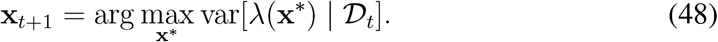

### 6.5 Results: adaptive stimulus selection for color tuning curves

We applied our method to the problem of estimating color tuning functions for V1 neurons recorded in awake, fixating monkeys under two paradigms. In the first paradigm, Gabor patches (Gaussian windowed sinusoidal gratings) were drifted across the V1 receptive field. The orientation, spatial frequency, and direction of drift in each Gabor patch was tailored to each neuron. The drifting Gabors modulated the long, medium, and short-wavelength sensitive cones in various degrees and ratios such that they produced a single target response from the neuron. In the second paradigm, Gaussian smoothed rectangular bars drifted across the receptive field. The length and width of the bar, as well as it direction of drift, was tailored to each cell and smoothed with a Gaussian window. Fig. 9 shows a schematic illustrating the stimulus space, the raw experimental data, and the adaptive stimulus stimulus selection protocol.

**Figure 9:**
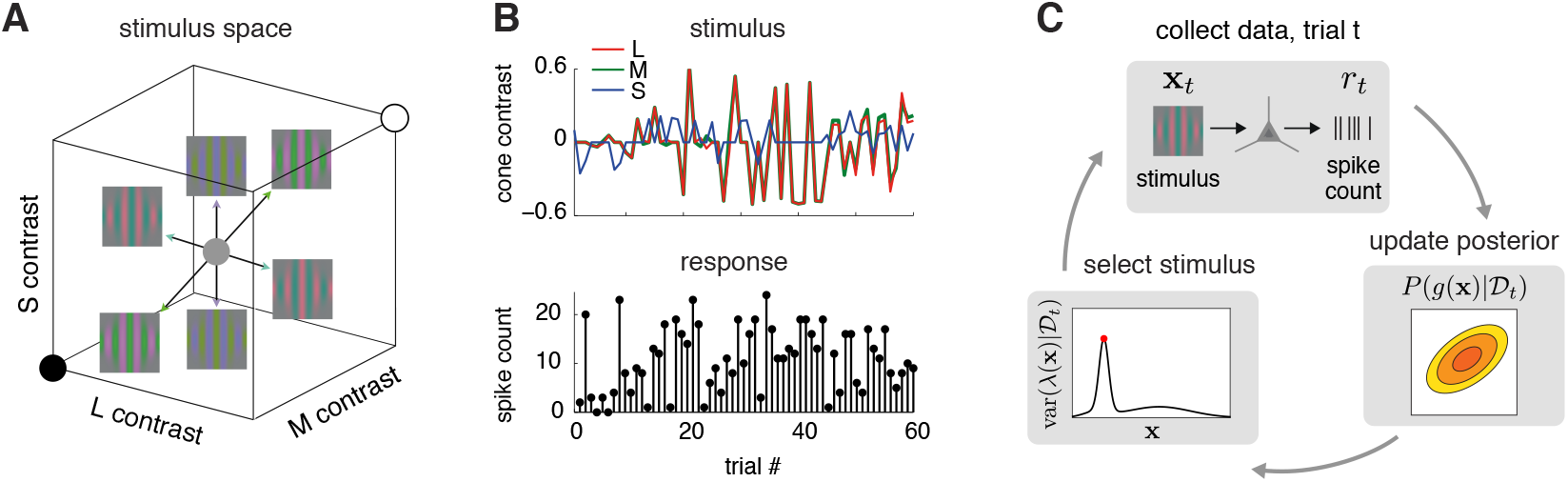
Schematic of stimulus space, data, and adapative stimulus selection method. **(A)** The stimulus on each trial was a drifting sinusoidal grating, aligned with the cell’s preferred orientation, with color defined by a pair of points symmetrically offset from zero in 3D cone-contrast space. The axes of this space are the inputs to long (L), medium (M), and short (S) wavelength cones, respectively, and the pair of points defining each stimulus indicate the grating’s spectral modulation amplitude and spectral orientation (i.e., where one point defines the color of the peaks and the other defines the color of the troughs of the sinusoid). The center of this space represents a neutral gray stimulus with no spectral modulation. **(B)** Example data for one cell: the stimulus on each trial is uniquely defined by three numbers (cone contrasts for L, M, and S in the upper-diagonal half-cube) and results in a spike count response measured over 667 ms. **(C)** Schematic of adaptive stimulus selection procedure. On each trial, we present stimulus **x**
_
*t*
_ and record spike response *r_t_
*. Next, we update the posterior over the tuning curve *g* using the full dataset collected so far, *D_t_
*. Lastly, we compute the posterior variance of the tuning curve, var(*λ*(**x**
^
*∗*
^) *D_t_
*), for a grid of points **x**
^
*∗*
^ spanning the stimulus space, and select the point with maximal variance as the stimulus for the next trial.

In interleaved trials, individual V1 neurons were tested with stimuli that were chosen either adaptively (using our method, Fig. 9C) or non-adaptively. In non-adaptive trials, stimulus selection was independent of responses. In adaptive trials, stimulus selection was based on the posterior variance of the firing rate map in the over-dispersed GP-Poisson model. Results showed that the adaptive method yielded faster convergence and more accurate firing map estimates than the non-adaptive method.

We used color-contrast-varying stimuli (either 2D or 3D), whose contrast levels are tuned by three values applied to L, M, and S cones. We measured responses to the stimuli by each cell from the primary visual cortex. As shown in Figure 10, the estimates of the Poisson model are much coarser than would be expected. As a result, the super-Poisson model achieved on average a four-times higher likelihood on test data, compared to the Poisson model.

**Figure 10:**
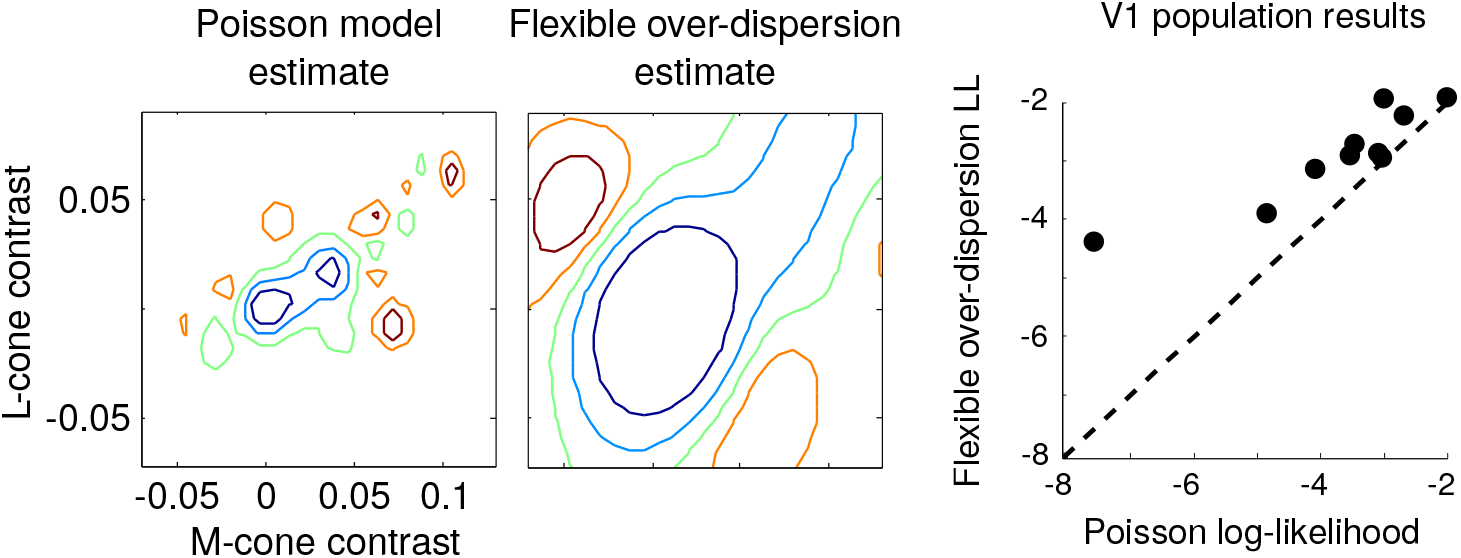
Comparison of tuning curve estimates under the GP-Poisson model and GP-flexible-over-dispersion model. **Left:** A 2D slice through the 3D color tuning curve for a V1 cell corresponding to the *L*-*M* plane (with *S* = 0) under Poisson (left) and flexible over-dispersion model (right). The flexible over-dispersion achieved a 25 times higher likelihood for the test dataset of 76 trials (763 total trials, 90%-10% split for cross validation), corresponding to 4.3% higher per-trial likelihood, and exhibited a smoother estimate (larger length scale *δ*, as observed in Fig. 8). **Right:** The test log-likelihood from 10 V1 cells, averaged over 10-fold cross validation. Each dot represents the log-likelihood of the data collected from each cell. On average, the flexible over-dispersion model achieved a 4 times higher test likelihood than the Poisson model, indicating its greater suitability for application to closed-loop experiments.

In the closed-loop design, we selected the stimulus that had the highest posterior variance of firing rate map in each trial. In the open-loop design, we sampled the stimuli uniformly from the polar grid. In Figure 11, we show the estimated tuning functions using all data collected from both paradigms, as well as estimates using a quarter of the data collected from each design. The estimates obtained by closed-loop design look more similar to the estimates obtained by using all the data. Finally, we computed the average prediction error using data collected from each paradigm. The closed-loop method achieved substantially lower prediction error and faster convergence than the open-loop method.

**Figure 11:**
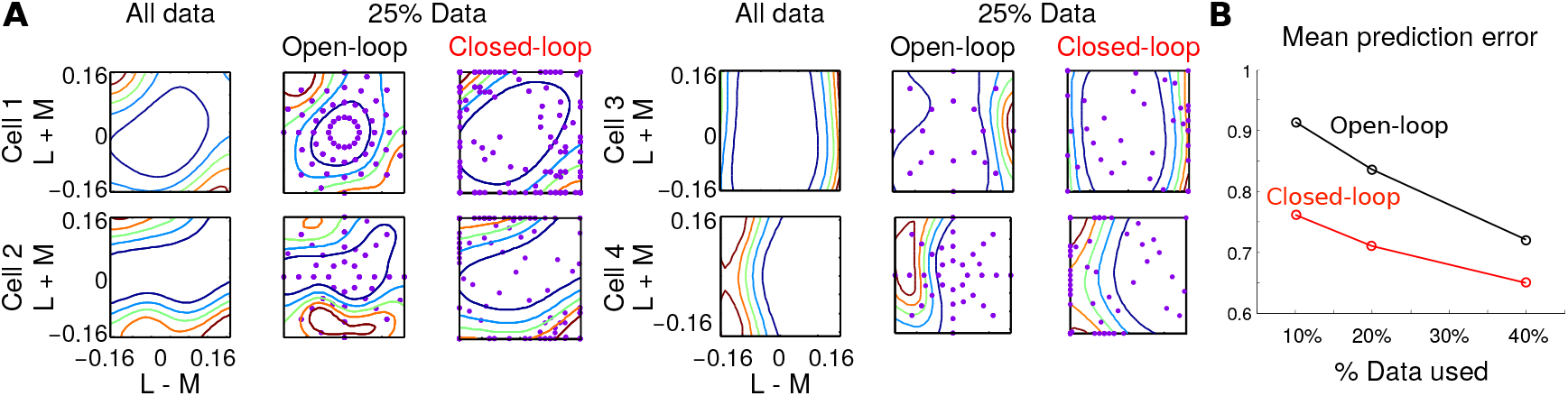
Closed-loop vs open-loop designs of neurophysiology experiments (a): Contour plot of the estimated firing rate maps of four different V1 cells. Purple dots indicate stimuli selected by each experimental paradigm. (b): Average prediction error as a function of the amount of data. The closed-loop method achieved 16% higher prediction accuracy compared to the open-loop method.

## 7 Discussion

We have presented here a flexible model for super-Poisson spike-count responses and applied the new model to closed-loop neurophysiology experiments. Our model, which introduces extra-Poisson variability via a Gaussian noise term added to the stimulus response level and passed through a nonlinearity, provides flexibility in accounting for neural over-dispersion and permits efficient model-fitting. We find that this model better fits individual mean-variance relationships for neurons in V1. Furthermore, we find that when used as a likelihood in lieu of the Poisson likelihood, our model can significantly improve closed-loop estimation of color tuning curves in V1.

### Relationship to previous work

A wide variety of other approaches to over-dispersed spike count distributions have been proposed in the recent literature. These include mixtures of Poisson distributions (Wiener & Richmond, 2003; Shidara et al., 2005) and the Twiddle distribution, which is characterized by a polynomial mean-variance relationship var[*r*] = *α*𝔼[*r*])^
*ρ*
^ for some *α* and *ρ* (Moshitch & Nelken, 2014; Koyama, 2015; Gershon et al., 1998). Although these models offer substantial improvements over models that specify a single Fano-factor, they are still restricted in the classes of over-dispersion modeled, and in some cases are more difficult to connect to mechanistic interpretations (i.e., how such counts might be generated). In contrast to these approaches, recent work on neural partitioning and negative binomial spike counts (Onken et al., 2009; Goris et al., 2014; Pillow & Scott, 2012a) instead seeks more descriptive, interpretable models of over-dispersion via hierarchical (or doubly stochastic) modeling.

Other methods to quantify neural over-dispersion rely on describing more general count distributions, such as the Bernoulli or Conway-Maxwell-Poisson (COM-Poisson) distributions (Zhu et al., 2017; Sellers et al., 2012), rather than layering on top of the Poisson distribution (Gao et al., 2015; Stevenson, 2016). These models may be limited by requiring many additional parameters (Gao et al., 2015), or may require that the mean-variance curves be monotonically increasing (Stevenson, 2016), reducing the model’s ability to explain neurons which exhibit higher variance at lower means. While completely replacing the Poisson distribution is satisfying in that the mean-variance relationship is no longer restricted at any stage by Poisson behavior, we find that layering additional variability inside the Poisson rate is more intuitive (Goris et al., 2014). Specifically, we find that such hierarchical models can isolates variables responsible for over-dispersion and allows mechanistic theories to attempt and explain these factors.

Our model can be considered as a significant generalization of the mixture-of-Poisson models (Wiener & Richmond, 2003; Shidara et al., 2005), as our model is essentially an infinite mixture of Poissons, as well as a generalization of prior work using latent hierarchical dispersion variables (Goris et al., 2014). In terms of the power-law models of (Moshitch & Nelken, 2014; Koyama, 2015), our model also includes as a subset any power-law where the exponent is between one and two (i.e. var[*r*] = *a*(𝔼[*r*])^
*ρ*
^ for 1*≤ ρ ≤*2). Although our model is incapable of achieving *ρ >* 2 or *ρ <* 1, the achievable range of *ρ* includes experimentally observed values in the literature, e.g. *ρ* ≈ 1.3 (Shadlen & Newsome, 1998) or 1.1 *≤ ρ ≤* 1.5 for cat auditory cortex (Moshitch & Nelken, 2014). Interestingly, however, our analysis demonstrates that power-law type characterizations might still be insufficient for assessing neural overdispersion. In particular, our model exploration indicates that a full discussion of the firing statistics should also include the full count distributions (see Fig. 7). Our results demonstrated that our model more accurately captures the variation in V1 neural responses, as compared to both traditional and more recent Poisson-based models.

### Interpretation and future directions

The origins of neural response variability are still a subject of debate (Renart & Machens, 2014; Masquelier, 2013), and we admit that our model does not attempt to explain how over-dispersion arises. Our model only implies that the unknown nuisance variable is well modeled by an addition of stimulus-dependent drive and Gaussian noise, followed by a nonlinearity and Poisson spike generation. In terms of the nonlinearity used, we found that the single-parameter “power soft-rectification” function was flexible enough to account for a large range of neural behavior, despite its reliance on a single parameter. The inference cost for the flexible overdispersion model therefore remains modest, and is moreover justified by the substantial gain in model expressibility (e.g., as shown by the AIC results in Fig. 6). Other nonlinearities are certainly possible in this framework, and the general properties we describe in Section 3 guarantee that, aside from perhaps requiring numerical integration, the model will be well behaved (i.e. the variance-mean curve will remain a proper function). For example, we found that the exponential non-linearity allowed for more computationally efficient calculations for the closed-loop inference procedure. We note that our model also does not attempt to account for the *under*-dispersion sometimes observed in certain neural circuits, such as retina or auditory cortex, that exhibit higher degrees of response regularity (DeWeese & Zador, 2002; Kara et al., 2000; Gur et al., 1997; Maimon & Assad, 2009). Comparable flexible models for under-dispersed spike counts therefore poses an important open problem for future work. As a final note, the increasing number of simultaneously recorded neurons also poses a future challenge of explaining the

While we focus here on explaining super-Poisson behavior of neural firing, we note that our model does not attempt to account for the *under*-dispersion sometimes observed in certain neural circuits, such as retina or auditory cortex, that exhibit higher degrees of response regularity (DeWeese & Zador, 2002; Kara et al., 2000; Gur et al., 1997; Maimon & Assad, 2009). Comparable flexible models for under-dispersed spike counts therefore poses an important open problem for future work. As a final note, our model focuses only on explaining the spiking process for a single neuron. The increasing number of simultaneously recorded neurons thus poses the future challenge of explaining correlations in over-dispersion between neurons (e.g. Goris et al. (2014)). Such population over-dispersion models would carry through the benefits of more accurate spike-count models that we observe here for single-neuron closed-loop experiments to cases where entire populations are analyzed simultaneously.

## Acknowledgments

We are grateful to Arnulf Graf and J. Anthony Movshon for V1 spike count data used for fitting and validating the flexible over-dispersion model. ASC was supported by the NIH NRSA Training Grant in Quantitative Neuroscience (T32MH065214). MP was supported by the Gatsby Charitable Foundation. JWP was supported by grants from the McKnight Foundation, Simons Collaboration on the Global Brain (SCGB AWD1004351) the NSF CAREER Award (IIS-1150186), and an NIMH grant (MH099611).

## Appendix A: Modulated Poisson and the NB model

Here we review the connection between the modulated Poisson model introduced by Goris et al. (2014) and the negative binomial model. The modulated Poisson model was introduced generally as having a Poisson rate *λ*(**x**) modulated by a stochastic gain *G* with a distribution satisfying 𝔼[*G*] = 1 and 
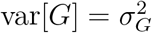
. The general mean-variance relationship of the marginal spike counts under this model was shown to always satisfy a quadratic relationship (i.e., 
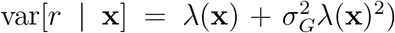
. However, fitting the model parameters to data required additional assumptions about the distribution of *G*. In particular, Goris *et al*. used a gamma distribution, which makes the integral over *G* analytically tractable, producing a negative binomial distribution over spike counts.

To see this connection formally, we can write the model, substituting *z* = *λ*(**x**) for the stimulus-dependent firing rate:

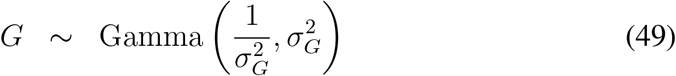

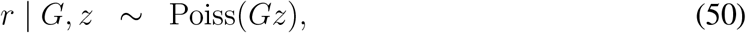

where 1/
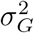
 and 
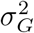
 correspond to the shape and scale parameters for the gamma distribution, which ensures 𝔼[*G*] = 1 and var[*G*] = 
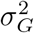
. The spike count *r* then has a negative binomial distribution:

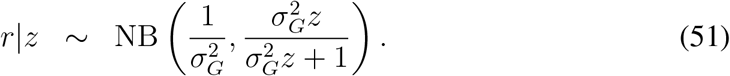

The derivation of the Negative Binomial PMF from a Poisson distribution with a Gamma prior can be obtained via the following straight-forward integration:

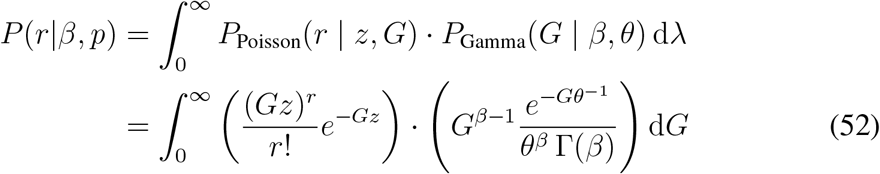

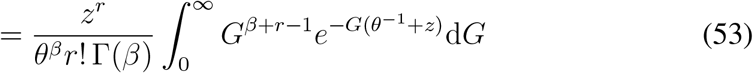

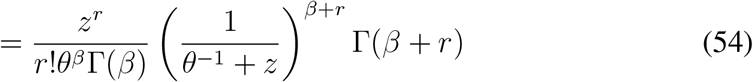

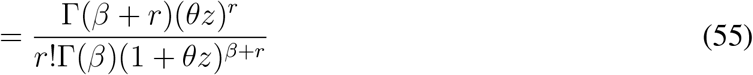

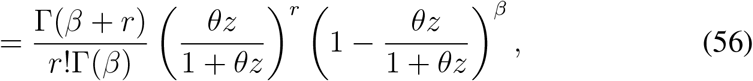

which for
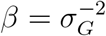
 and
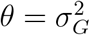
 results in the above NB distribution.

## Appendix B: Parameter optimization using Laplace approximation

The general method we present for estimating the parameters of the nonlinearity that modulates neural spiking is outlined in Algorithm 1. For a particular nonlinearity (i.e. the soft-rectification raised to a power in Equation (13)), these steps can be written more explicitly. Here we fit data using both the exponential nonlinearity, for which the fitting algorithm is provided in Algorithm 3, and the power-soft-rectification nonlinearity, corresponding to the model fitting in Algorithm 2). Note that the estimation of and the gradient descent steps for *θ* and *z* can be accomplished analytically, however the estimation of *n*
_0_ would typically need to be calculated numerically.

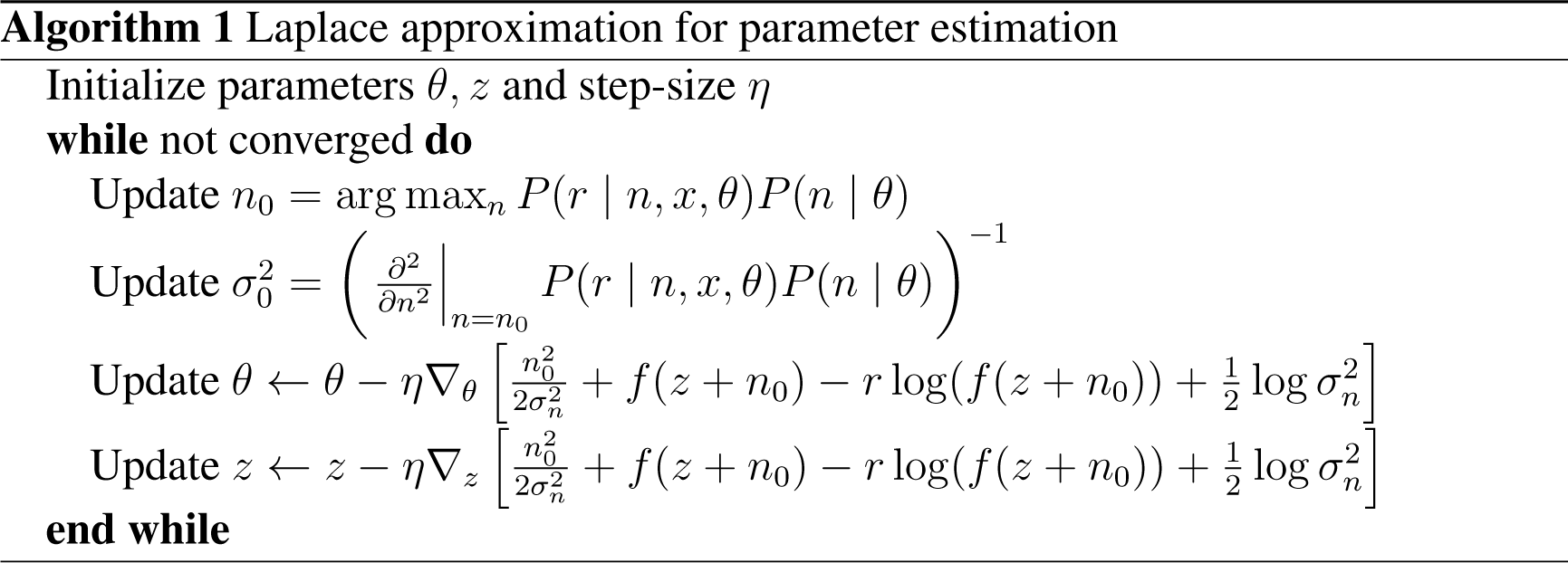

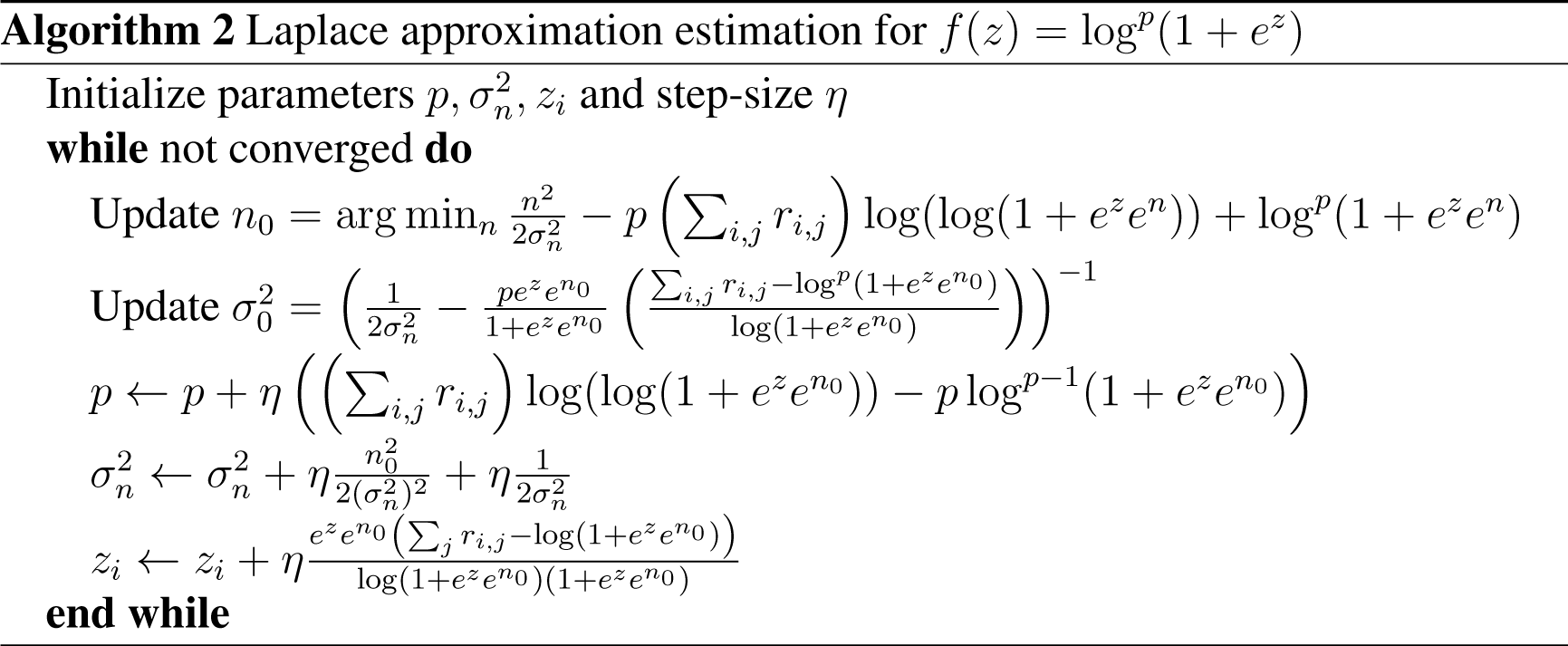

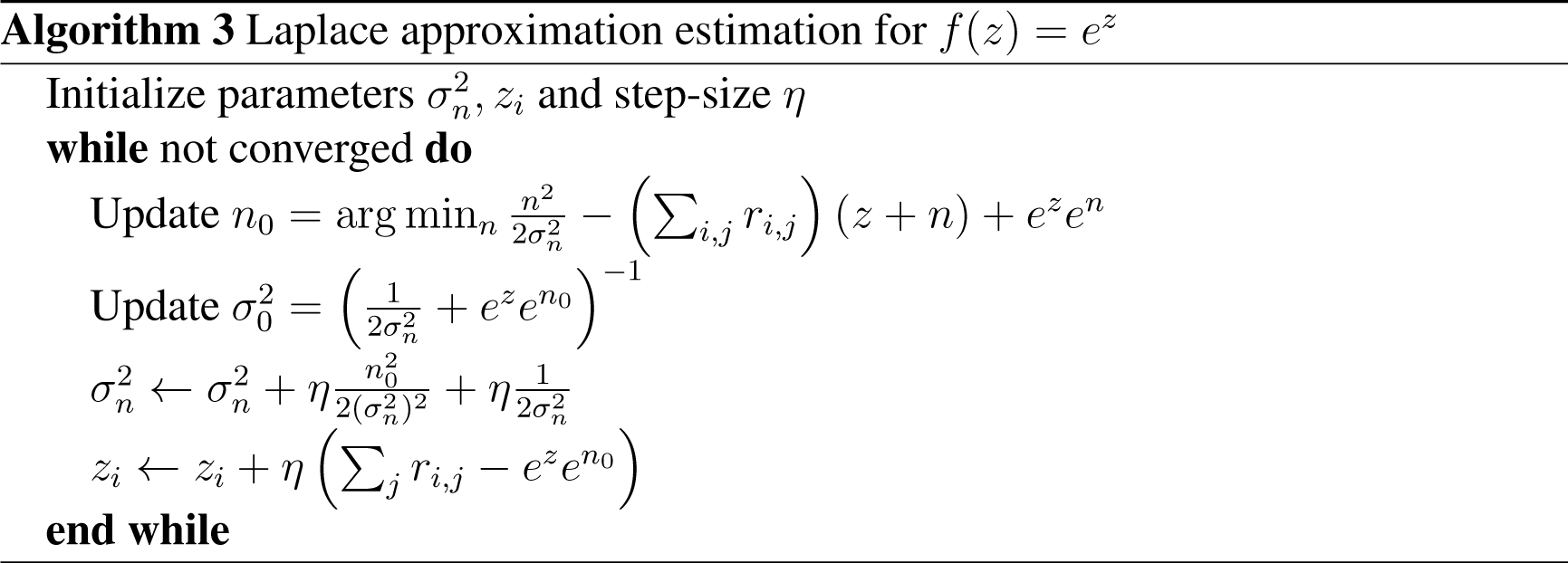

## Footnotes

1 Note that in contrast to (Goris et al., 2014), we define *λ*(**x**) in units of spikes-per-bin, removing the bin-size Δ from the ensuing equations.

## References

Archer, E. W., Koster, U., Pillow, J. W., & Macke, J. H. (2014). Low-dimensional models of neural population activity in sensory cortical circuits. In Advances in Neural Information Processing Systems 27, Z. Ghahramani, M. Welling, C. Cortes, N. Lawrence, & K. Weinberger, eds. (Curran Associates, Inc.), pp. 343–351.

Baddeley, R., Abbott, L. F., Booth, M. C., Sengpiel, F., Freeman, T., Wakeman, E. A., & Rolls, E. T. (1997). Responses of neurons in primary and inferior temporal visual cortices to natural scenes. Proceedings of the Royal Society of London B: Biological Sciences, 264, 1775–1783.

Barberini, C., Horwitz, G., & Newsome, W. (2001). A comparison of spiking statistics in motion sensing neurones of flies and monkeys. In Motion Vision. (Springer), pp. 307–320.

Benda, J., Gollisch, T., Machens, C. K., & Herz, A. V. (2007). From response to stimulus: adaptive sampling in sensory physiology. Curr. Opin. Neurobiol., 17, 430–436.

Bernardo, J. M. (1979). Expected information as expected utility. The Annals of Statistics, 7, 686–690.

Bölinger, D. & Gollisch, T. (2012). Closed-loop measurements of iso-response stimuli reveal dynamic nonlinear stimulus integration in the retina. Neuron, 73, 333–346.

Brillinger, D. R. (1988). Maximum likelihood analysis of spike trains of interacting nerve cells. Biological cybernetics, 59, 189–200.

Buesing, L., Macke, J., & Sahani, M. (2012). Spectral learning of linear dynamics from generalised-linear observations with application to neural population data. In Advances in Neural Information Processing Systems 25, pp. 1691–1699.

Buracas, G., Zador, A., DeWeese, M., & Albright, T. (1998). Efficient discrimination of temporal patterns by motion-sensitive neurons in primate visual cortex. Neuron, 5, 959–969.

Carandini, M. (2004). Amplification of trial-to-trial response variability by neurons in visual cortex. PLoS biology, 2, e264.

Chaloner, K. & Verdinelli, I. (1995). Bayesian experimental design: a review. Statistical Science, 10, 273–304.

Chichilnisky, E. J. (2001). A simple white noise analysis of neuronal light responses. Network: Computation in Neural Systems, 12, 199–213.

Cohn, D. A., Ghahramani, Z., & Jordan, M. I. (1996). Active learning with statistical models. J. Artif. Intell. Res. (JAIR), 4, 129–145.

DeWeese, M. & Zador, A. M. (2002). Binary coding in auditory cortex. In NIPS.

DiMattina, C. & Zhang, K. (2011). Active data collection for efficient estimation and comparison of nonlinear neural models. Neural computation, 23, 2242–2288.

DiMattina, C. & Zhang, K. (2013). Adaptive stimulus optimization for sensory systems neuroscience. Frontiers in neural circuits, 7.

Ecker, A. S., Denfield, G. H., Bethge, M., & Tolias, A. S. (2016). On the structure of neuronal population activity under fluctuations in attentional state. The Journal of Neuroscience, 36, 1775–1789.

Eden, U. & Kramer, M. (2010). Drawing inferences from fano factor calculations. Journal of neuroscience methods, 190, 149–152.

Gao, Y., Archer, E. W., Paninski, L., & Cunningham, J. P. (2016). Linear dynamical neural population models through nonlinear embeddings. In Advances in Neural Information Processing Systems, pp. 163–171.

Gao, Y., Busing, L., Shenoy, K., & Cunningham, J. (2015). High-dimensional neural spike train analysis with generalized count linear dynamical systems. In Advances in Neural Information Processing Systems, pp. 2044–2052.

Geisler, W. S. & Albrecht, D. G. (1997). Visual cortex neurons in monkeys and cats: detection, discrimination, and identification. Vis Neurosci, 14, 897–919.

Gershon, E. D., Wiener, M. C., Latham, P. E., & Richmond, B. J. (1998). Coding strategies in monkey v1 and inferior temporal cortices. J Neurophysiol, 79, 1135–1144.

Goris, R. L. T., Movshon, J. A., & Simoncelli, E. P. (2014). Partitioning neuronal variability. Nat Neurosci, 17, 858–865.

Graf, A. B., Kohn, A., Jazayeri, M., & Movshon, J. A. (2011). Decoding the activity of neuronal populations in macaque primary visual cortex. Nature neuroscience, 14, 239–245.

Gur, M., Beylin, A., & Snodderly, D. M. (1997). Response variability of neurons in primary visual cortex (v1) of alert monkeys. J Neurosci, 17, 2914–2920.

Kara, P., Reinagel, P., & Reid, R. C. (2000). Low response variability in simultaneously recorded retinal, thalamic, and cortical neurons. Neuron, 27, 636–646.

Koyama, S. (2015). On the spike train variability characterized by variance-to-mean power relationship. Neural computation, 27, 1530–1548.

Lewi, J., Butera, R., & Paninski, L. (2009). Sequential optimal design of neurophysiology experiments. Neural Computation, 21, 619–687.

Linderman, S., Adams, R. P., & Pillow, J. W. (2016). Bayesian latent structure discovery from multi-neuron recordings. In Advances in Neural Information Processing Systems 29, D. D. Lee, M. Sugiyama, U. V. Luxburg, I. Guyon, & R. Garnett, eds. (Curran Associates, Inc.), pp. 2002–2010.

Lindley, D. (1956). On a measure of the information provided an experiment. Ann. Math. Statist., 27, 986–1005.

MacKay, D. (1992). Information-based objective functions for active data selection. Neural Computation, 4, 589–603.

Macke, J. H., Buesing, L., Cunningham, J. P., Byron, M. Y., Shenoy, K. V., & Sahani, M. (2011). Empirical models of spiking in neural populations. In Advances in neural information processing systems, vol. 24, pp. 1350–1358.

Maimon, G. & Assad, J. A. (2009). Beyond poisson: increased spike-time regularity across primate parietal cortex. Neuron, 62, 426–440.

Masquelier, T. (2013). Neural variability, or lack thereof. Frontiers in computational neuroscience, 7, 7.

Moshitch, D. & Nelken, I. (2014). Using tweedie distributions for fitting spike count data. Journal of neuroscience methods, 225, 13–28.

Nelder, J. & Baker, R. (1972). Generalized linear models. Encyclopedia of statistical sciences.

Onken, A., Grünewälder, S., Munk, M. H., & Obermayer, K. (2009). Analyzing shortterm noise dependencies of spike-counts in macaque prefrontal cortex using copulas and the flashlight transformation. PLoS Comput Biol, 5, e1000577.

Paninski, L. (2004). Maximum likelihood estimation of cascade point-process neural encoding models. Network: Computation in Neural Systems, 15, 243–262.

Paninski, L. (2005). Asymptotic theory of information-theoretic experimental design. Neural Computation, 17, 1480–1507.

Paninski, L., Pillow, J. W., & Lewi, J. (2007). Statistical models for neural encoding, decoding, and optimal stimulus design. In Computational Neuroscience: Theoretical Insights Into Brain Function, P. Cisek, T. Drew, & J. Kalaska, eds., Progress in Brain Research. (Elsevier), pp. 493–507.

Park, M. & Pillow, J. W. (2012). Bayesian active learning with localized priors for fast receptive field characterization. In Advances in Neural Information Processing Systems 25, P. Bartlett, F. Pereira, C. Burges, L. Bottou, & K. Weinberger, eds., pp. 2357–2365.

Park, M., Weller, J. P., Horwitz, G. D., & Pillow, J. W. (2014). Bayesian active learning of neural firing rate maps with transformed gaussian process priors. Neural Computation, 26, 1519–1541.

Pillow, J. & Scott, J. (2012a). Fully bayesian inference for neural models with negative-binomial spiking. In Advances in Neural Information Processing Systems 25, P. Bartlett, F. Pereira, C. Burges, L. Bottou, & K. Weinberger, eds., pp. 1907–1915.

Pillow, J. W. & Park, M. (2016). Adaptive bayesian methods for closed-loop neurophysiology. In Closed Loop Neuroscience, A. El Hady, ed. (Elsevier), pp. 3–18.

Pillow, J. W. & Scott, J. G. (2012b). Fully bayesian inference for neural models with negative-binomial spiking. In NIPS, pp. 1907–1915.

Rabinowitz, N. C., Goris, R. L., Cohen, M., & Simoncelli, E. (2015). Attention stabilizes the shared gain of v4 populations. eLife.

Rad, K. R. & Paninski, L. (2010). Efficient, adaptive estimation of two-dimensional firing rate surfaces via gaussian process methods. Network: Computation in Neural Systems, 21, 142–168.

Rasmussen, C. & Williams, C. (2006). Gaussian Processes for Machine Learning. (MIT Press).

Renart, A. & Machens, C. K. (2014). Variability in neural activity and behavior. Current opinion in neurobiology, 25, 211–220.

Sellers, K. F., Borle, S., & Shmueli, G. (2012). The com-poisson model for count data: a survey of methods and applications. Applied Stochastic Models in Business and Industry, 28, 104–116.

Shadlen, M. & Newsome, W. (1998). The variable discharge of cortical neurons: implications for connectivity, computation, and information coding. Journal of Neuroscience, 18, 3870–3896.

Shidara, M., Mizuhiki, T., & Richmond, B. J. (2005). Neuronal firing in anterior cingulate neurons changes modes across trials in single states of multitrial reward schedules. Experimental brain research, 163, 242–245.

Simoncelli, E. P., Pillow, J. W., Paninski, L., & Schwartz, O. (2004). Characterization of neural responses with stochastic stimuli. In The Cognitive Neurosciences, III, M. Gazzaniga, ed. (Cambridge, MA: MIT Press), chap. 23, pp. 327–338.

Stevenson, I. H. (2016). Flexible models for spike count data with both over-and underdispersion. Journal of computational neuroscience, 41, 29–43.

Tolhurst, D. J., Movshon, J. A., & Dean, A. F. (1983). The statistical reliability of signals in single neurons in cat and monkey visual cortex. Vision Res, 23, 775–785.

Wiener, M. C. & Richmond, B. J. (2003). Decoding spike trains instant by instant using order statistics and the mixture-of-poissons model. Journal of Neuroscience, 23, 2394–2406.

Zhao, Y. & Park, I. M. (2017). Variational latent gaussian process for recovering singletrial dynamics from population spike trains. Neural Computation.

Zhu, L., Sellers, K. F., Morris, D. S., & Shmueli, G. (2017). Bridging the gap: a generalized stochastic process for count data. The American Statistician, 71, 71–80.

